# Abnormal neuronal and synaptic morphology in Down syndrome brains reproduces in human isogenic cellular models

**DOI:** 10.64898/2026.01.10.698781

**Authors:** Ante Plećaš, Gillian Gough, Danka Grčević, Hanna Jackowiak, Kinga Skieresz-Szewczyk, Matija Horaček, Iva Šimunić, Željka Krsnik, Dinko Mitrečić, Aoife Murray, Dean Nižetić, Ivan Alić

## Abstract

Down syndrome (DS) is the most common autosomal aneuploidy compatible with postnatal survival and is caused by full or partial trisomy of chromosome 21. In this study, human isogenic induced pluripotent stem cells (iPSCs) were differentiated into 2D neurons and cortico-striatal assembloids (hCSAs). Abnormal neuronal morphology was observed: trisomic (T21) neurons formed aggregates of cell bodies interconnected with thick neurite bundles radiating outwards, connected and overlapped with other neurons in between the bundles while disomic (D21) neurons were evenly distributed across the surface and formed strong neuronal networks. Detailed analysis revealed significantly shorter neurites with larger diameter, fewer branches, and fewer terminal points in T21 neurons compared to D21 neurons in both 2D cultures and hCSAs. We observed similar phenotypes in foetal and postnatal human brain tissue. Furthermore, we observed abnormal mitochondrial morphology with an excess of some mitochondrial proteins (AIF, TOMM20), abnormal synaptic morphology and significantly lower expression of both presynaptic and postsynaptic markers (SYN-1, PSD95 and GEPH) throughout *in vitro* differentiation of T21 neurons. Finally, our data showed that T21 spheroids were significantly smaller throughout *in vitro* differentiation compared to D21 spheroids. T21 spheroids also exhibited significantly higher expression of neural stem cells (SOX2), significantly lower expression of proliferating cells (Ki67) and significantly higher expression of apoptotic cells (CCaspase-3). Overall, our study demonstrates that T21 leads to abnormal neuronal morphology in both 2D neurons and hCSAs, consistent with observations in the human brain.

## INTRODUCTION

Down Syndrome (DS) is the most common autosomal aneuploidy compatible with postnatal survival, caused by full or partial trisomy of human chromosome 21 (HSA21). It affects over 6 million people worldwide and is characterised by alterations in most organ systems, in particular the cardiovascular, musculoskeletal and nervous system [1]. Moreover, DS is the most common genetic cause of intellectual disability and early-onset Alzheimer’s disease (AD) [2]. In addition to full or partial trisomy of chromosome 21, mosaic DS [3–5] is another variant of DS, used as a genetic tool in our research.

Neurofilaments are composed of five intermediate filaments [6] which can affect synapse density, morphology and functioning [6–8] particularly N-methyl-D-aspartate (NMDA) transmission in excitatory neurons [9, 10]. Previous studies using the Ts65Dn mouse model have shown that NMDA synaptic transmission is particularly disrupted in DS [11]. Furthermore, neurofilaments are involved in microtubule organisation, intracellular transport of molecules and organelles [12] as well as outgrowth and formation of both dendrites and axons [13, 14]. They also play a critical role in maturation and function of dendritic spines as well as in the regulation of glutamatergic and dopaminergic neurotransmission [6, 15, 16], which is highly relevant for our study, as our 2D cultures contain over 93% glutamatergic neurons [17]. Since mitochondria are involved in synaptic function [18, 19], and their biology and cellular distribution are linked to neurite branching and termini formation [20–22], we hypothesized that mitochondrial dysfunction and abnormal distribution in DS impair neurite branching, a defect previously documented in DS foetal neuronal cultures [23]. Importantly, neurofilament dysfunction and/or aggregation has been implicated in AD [24] and other neurodegenerative disorders including DS. Taken together, these findings encouraged us to study neuronal morphology using isogenic human induced pluripotent stem cells (iPSCs) [3] as a model.

Over the last decade, several isogenic iPSC [3, 25–28] models have been reported. Although some morphological features in T21 cells have been reported in embryonic stem cells (ESCs) [29], primary DS neurons [23] and iPSC-derived GABAergic neurons [27], a detailed analysis of neuronal morphology caused by trisomy 21 is still missing in literature. Based on available literature and our own observations we asked whether T21 causes abnormal neuronal morphology in DS. To address this fundamental question, we used human isogenic iPSCs [3, 30] and differentiated them into 2D neurons and human cortico-striatal assembloids (hCSAs) [31, 32], composed of human cortical spheroids (hCSs) and human striatal spheroids (hStrSs). Additionally, in hStrSs 10% of GFP labelled iPSCs [30] were added to visualize neuronal morphology in 3D. Finally, our *in vitro* data were compared to the human brain [33].

Our results highlight that the additional copy of HSA21 in individuals with DS causes abnormal neuronal morphology; T21 neurons formed aggregates of cell bodies interconnected by thick neurite bundles radiated outwards, connected and overlapped with other neurons, while D21 neurons were evenly distributed across the surface forming strong neuronal networks. Trisomic neurons showed shorter and thicker neurites, with smaller number of branching and smaller number of terminal points. Moreover, we observed abnormal T21 mitochondrial morphology in our system, while the synaptic development and maturation were delayed in T21 neurons. In addition to altered neuronal morphology, our study also revealed a slower differentiation of T21 cells, delayed maturation and increased cell death. Overall, our study demonstrates that human-derived iPSCs and various *in vitro* models reliably recapitulate key events occurring in the human brain, providing a powerful alternative to human tissue, which is often inaccessible.

## RESULTS

### Isogenic T21 neurons showed abnormal morphology

The main goal of this study was to analyse neuronal morphology using an isogenic model of DS in 2D neurons and hCSAs, in comparison to the neuronal morphology in human brain in individuals with DS. The study was conducted using eight clones of iPSCs, which were validated for chromosome number (Supplementary Fig. 1a) and pluripotency (Supplementary Fig. 1b). Neural stem cells (NSCs) (Supplementary Fig. 1c) were then derived from these iPSCs and differentiated into 2D neurons (Fig. 1a). Throughout *in vitro* differentiation, the isogenic neurons exhibited following morphological features: T21 neurons formed aggregates of cell bodies interconnected with thick neurite bundles radiating outwards, connected and overlapped with other neurons in between the bundles, while D21 neurons were evenly distributed across the surface forming strong neuronal networks (Fig. 1a). At day *in vitro* 14 (DIV14), isogenic T21 cells showed significantly more SOX2 positive cells and 4-fold fewer DCX positive neurons, suggesting delayed neuronal differentiation in T21 cells (Fig. 1b-d). Although morphological differences between T21 and D21 cultures were observed, TUBB3 expression analysed by Western blotting (WB) showed increase by DIV60 in both, T21 and D21 neurons and reached comparable levels (Fig. 1e-f) as well as *MAP2* analysed by qPCR (Fig. 1g). Interestingly, in our conditions over 93% neurons were excitatory [17], VGLUT1 positive neurons (Fig. 1h-i) with a limited percentage of VGAT positive neurons (Fig. 1h). Nevertheless, T21 cultures showed significantly less MAP2 positive neurons per field of view by DIV60 analysed by Imaris 9.9.1 software (Fig. 1j-k). Next, we asked whether astrocytes were present in our isogenic system. In 2D cultures, we observed a limited number of astrocytes at DIV40, which showed a 6.6-fold increase in GFAP surface area compared to D21 cultures, with no change in MAP2 surface area. By DIV60, that difference was reduced, but the MAP2 surface remained lower and GFAP surface area increased in T21 vs D21 cells (Fig. 1j-l).

**Fig. 1.**
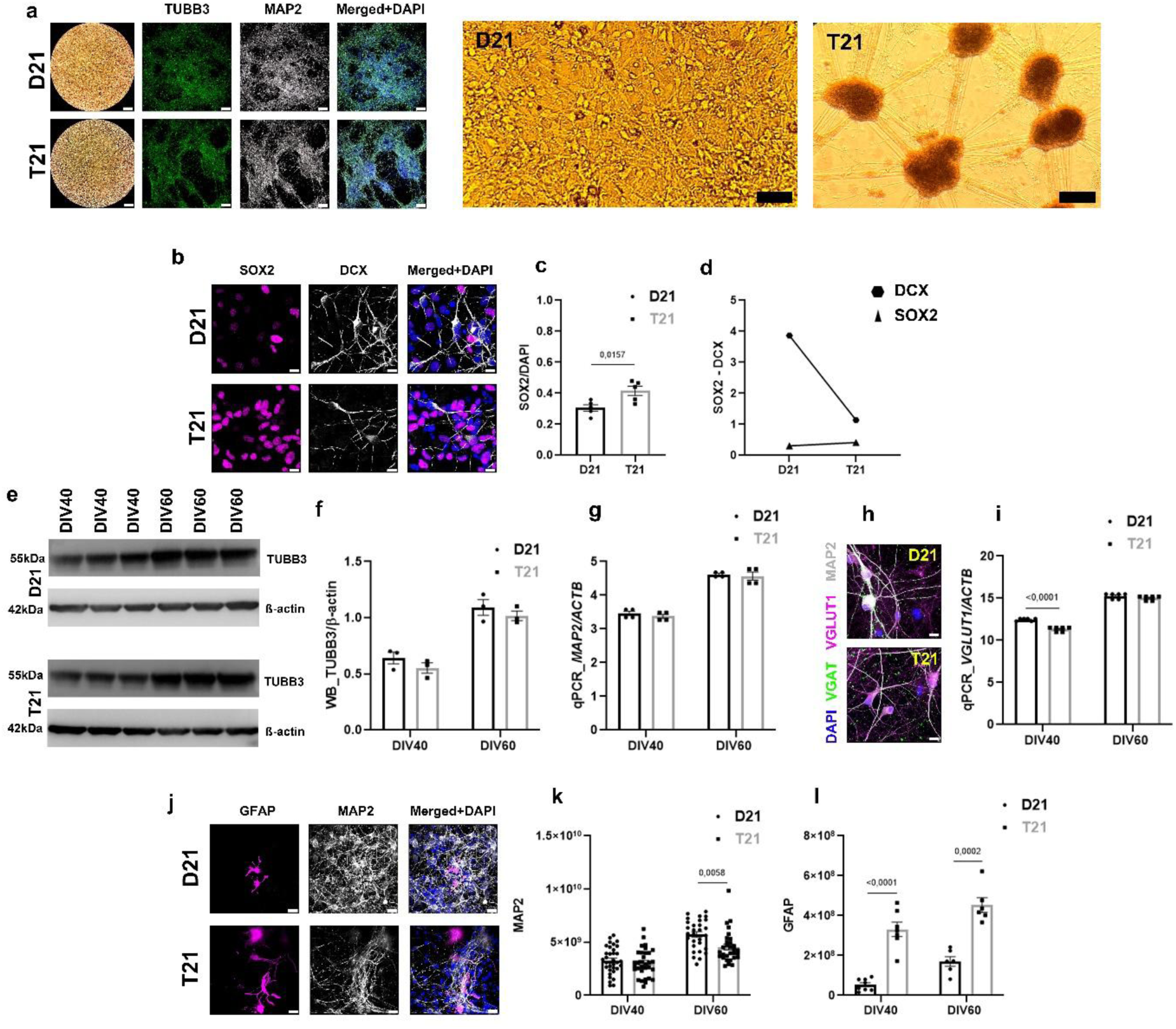
***In vitro* differentiation of iPSCs to 2D isogenic neurons**. **(a)** Representative brightfield images of isogenic neurons at DIV14 (left panel) and at DIV40 (right panel) showed abnormal T21 neuronal distribution in 2D. Representative confocal images of isogenic neurons stained with TUBB3 (green), MAP2 (white) and DAPI (blue) at DIV30. Scale bar: 100 µm. **(b)** Representative confocal images of isogenic cells stained with SOX2 (magenta), DCX (white) and DAPI (blue) at DIV14. Scale bar: 10 µm. **(c)** T21 cells showed significantly more SOX2 normalised by DAPI compared to the D21 cells. **(d)** Comparison of SOX2 and DCX expression showed significantly more SOX2 (triangle) positive NSCs and significantly fewer DCX (hexagon) positive neurons in T21 cells at DIV14. The y-axis (c, d) represents expression of SOX2 and/or DCX normalised by DAPI, while x-axis represents genotype. **(e)** Representative blots of TUBB3/ß-actin throughout *in vitro* differentiation. **(f)** WB analysis of TUBB3/ß-actin showed comparable protein expression at DIV40 and DIV60. The y-axis represents expression of TUBB3 normalised by β-actin, while x-axis represents DIV40 and DIV60. **(g)** Analysis of *MAP2/ACTB* throughout *in vitro* differentiation showed comparable gene expression at DIV40 and DIV60. The y-axis represents gene expression of *MAP2* normalised by *ACTB*, while x-axis represents DIV40 and DIV60. **(h)** Representative confocal images of isogenic neurons stained with VGAT (green), VGLUT1 (magenta), MAP2 (white) and DAPI (blue) at DIV60. Scale bar: 10 µm. **(i)** Analysis of *VGLUT1/ACTB* throughout *in vitro* differentiation showed the lower gene expression of *VGLUT1* in T21 neurons. The y-axis represents gene expression of *VGLUT1* normalised by *ACTB*, while x-axis represents DIV40 and DIV60. **(j)** Representative confocal images of isogenic neurons stained with GFAP (magenta), MAP2 (white) and DAPI (blue) at DIV60. Scale bar: 10 µm. **(k)** Total surface of MAP2 per field of view showed significantly fewer T21 neurons at DIV60. The y-axis represents total fluorescent surface of MAP2 per field of view, while x-axis represents DIV40 and DIV60. **(l)** Total surface of GFAP per field of view showed significantly more T21 astrocytes throughout differentiation. The y-axis represents total fluorescent surface of GFAP per field of view, while x-axis represents DIV40 and DIV60. Graphs represent means ± SEM.

Furthermore, isogenic hCSAs were grown from the same iPSCs. These hCSAs were generated by fusing hCSs and hStrSs. hCSAs were stained with MAP2 and GFAP and visualised by light-sheet microscopy (Fig. 2a). At DIV0, 10% of isogenic GFP labelled cells were admixed to the hStrSs-destined iPSCs, which enable easier cell tracing within the hCSAs (Fig. 2b). Importantly, the percentage of GFP positive cells remained stable throughout differentiation of hCSAs; by DIV70 3.37% GFP positive cells in D21 and 4.61% GFP positive cells in T21 hCSAs were detected by flow cytometry after dissociation of hCSAs (Fig. 2b). Moreover, that result was confirmed by WB throughout *in vitro* differentiation of hCSAs from DIV70 until DIV130 and showed similar data in isogenic hStrSs, while in control hCSs there was no GFP signal at DIV100 (Fig. 2c-d). Similarly, as in 2D neurons, TUBB3 expression in isogenic hCSAs remained at comparable levels throughout *in vitro* differentiation (Fig. 2e-f) indicating that our isogenic model consists predominantly of mature neurons. Although *MAP2* gene expression showed similar pattern and level of expression in both T21 and D21 hCSAs, significantly lower *MAP2* expression were observed in T21 hCSAs at DIV70 and DIV100 (Fig. 2h).

**Fig. 2.**
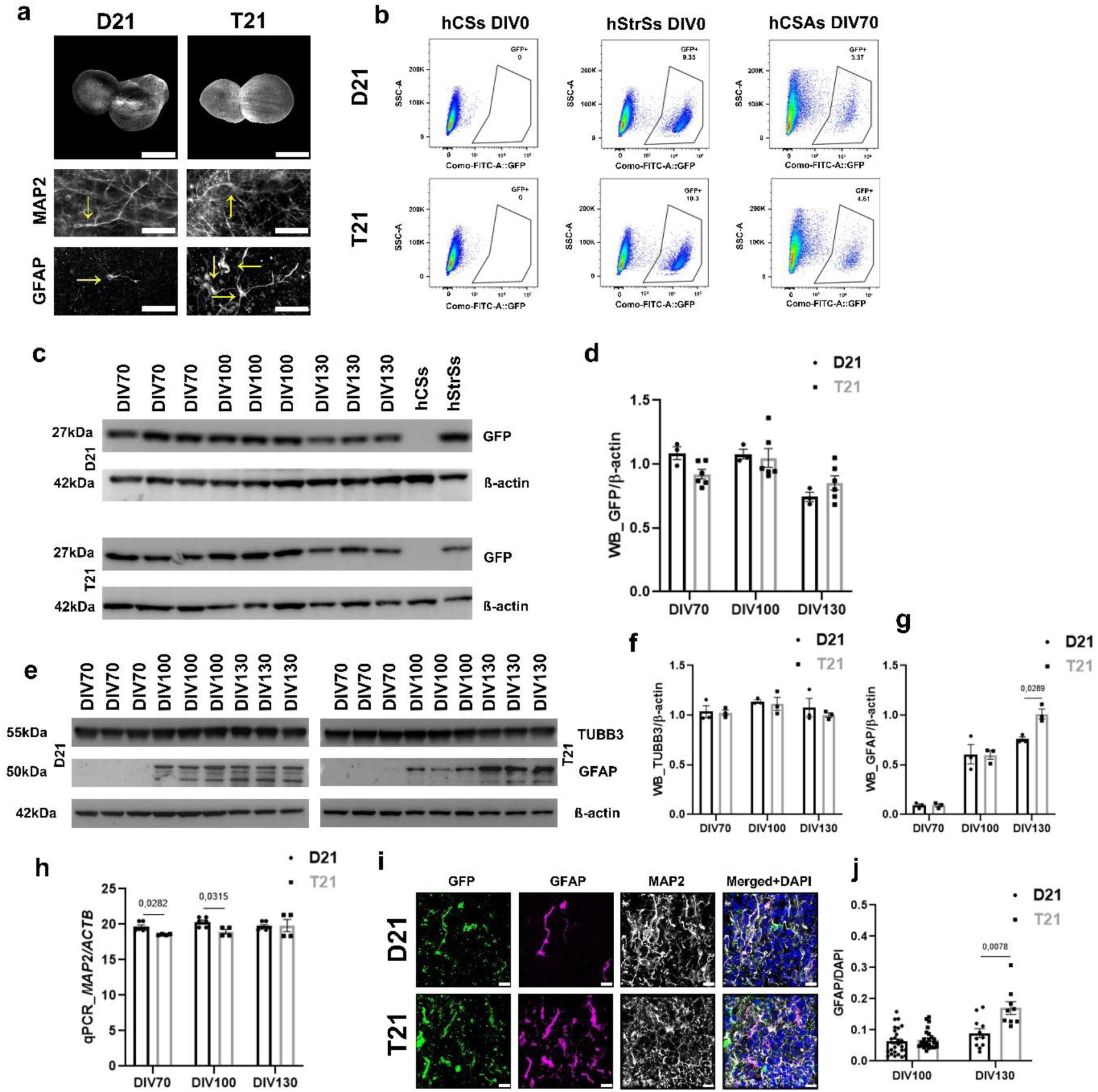
***In vitro* differentiation of isogenic hCSAs**. **(a)** Representative light-sheet images of cleared isogenic hCSAs at DIV100 stained with MAP2 and GFAP (top row). Scale bar: 500 µm. T21 hCSAs showed abnormal morphology of MAP2 positive neurons (middle row) and more GFAP positive astrocytes (bottom row). Scale bar: 50 µm. **(b)** Analysis of GFP expression by flow cytometry: representative plots of hCSs-destined iPSCs without GFP positive cells (left column), representative plots of hStrSs-destined iPSCs with 10% GFP positive cells at DIV0 (middle column) and representative plots of hCSAs showed 3.37% GFP positive cells in D21 and 4.61% GFP positive cells in T21 hCSAs at DIV70 (right column). The y-axis (SSC-A) represents signal area on side scatter (SSC) detector, while x-axis represents fluorescent intensity of GFP signal area. **(c)** Representative blots of GFP/ß-actin throughout *in vitro* differentiation. **(d)** WB analysis of GFP/ß-actin showed comparable protein expression, except in hCSs (control sample) at DIV100. The y-axis represents expression of GFP normalised by β-actin, while x-axis represents DIV70, DIV100 and DIV130. **(e)** Representative blots of TUBB3/ß-actin and GFAP/ß-actin throughout *in vitro* differentiation. **(f)** WB analysis of TUBB3/ß-actin showed comparable expression of TUBB3 throughout *in vitro* differentiation. The y-axis represents expression of TUBB3 normalised by β-actin, while x-axis represents DIV70, DIV100 and DIV130. **(g)** WB analysis of GFAP/ß-actin showed significantly more astrocytes in T21 hCSAs at DIV130. **(h)** Analysis of *MAP2/ACTB* throughout *in vitro* differentiation showed comparable gene expression in both D21 and T21 hCSAs with significantly lower expression in T21 hCSAs at DIV70 and DIV100. The y-axis represents expression of *MAP2* normalised by *ACTB*, while x-axis represents DIV70, DIV100 and DIV130. **(i)** Representative confocal images of isogenic hCSAs with GFP positive cells (green), stained with GFAP (magenta), MAP2 (white) and DAPI (blue) at DIV130. Scale bar: 10 µm. **(j)** The first astrocytes were observed at DIV100 and differentiation by DIV130 significantly more astrocytes normalised by DAPI, were observed in T21 hCSAs. The y-axis represents total fluorescent surface of GFAP normalised by DAPI, while x-axis represents DIV100 and DIV130. Graphs represent means ± SEM.

In addition to pan-neuronal markers, region-specific neuronal fate was assessed by analysing CTIP2, a marker of the deeper cortical layer (layer V) and SATB2, a marker of the superficial cortical layers (layers II/III). Although both markers were cortical, we found some striatal expression of both markers in isogenic hCSAs. Notably, CTIP2 expression was significantly higher in T21 neurons, in both hCSs and hStrSs (Supplementary Fig. 2) suggesting some skewing of the subtype fate during the development of T21 cortical layers. However, there was no difference in SATB2 expression between T21 and D21 hCSAs (Supplementary Fig. 2).

Furthermore, in the hCSAs, astrocytes started to be detectable at DIV100 (Fig. 2a, e, g). By DIV130, T21 hCSAs showed a 1.3-fold higher astrocyte presence compared to D21 hCSAs (Fig. 2i-j). Although our media composition directs NSCs primarily toward neuronal differentiation, we observed a certain percentage of astrocytes in both 2D and 3D systems. This percentage was significantly higher in T21 cells, suggesting a slight shift in the oligopotency of T21 NSCs toward an astrocytic cell fate.

Importantly, our isogenic cells did not show any inter-clonal variations between D21 (C3, C7, C9, and C3GFP) and T21 cells (C5, C6, C13, and C5GFP); therefore, in all figures, we pooled the results according to genotype.

### T21 neurons showed lager diameter and shorter neurites

In order to describe and analyse observed neuronal morphology, quantitative analysis of confocal images was performed by Imaris 9.9.1 software. Isogenic T21 neurons (Fig. 3a) showed significantly larger diameter of MAP2 positive dendrites throughout *in vitro* differentiation (Fig. 3b). At the same time, T21 dendrites were significantly shorter (Fig. 3c) with significantly fewer branches (Fig. 3d) compared to D21 neurons. We then asked whether the same phenotypes were present in hCSAs neurons (Fig. 3e). Representative 20,000x TEM images of isogenic dendrites showed the same pattern: the T21 dendrite measured 3.0 µm in diameter, whereas the D21 dendrite measured 2.6 µm. Imaris analysis was performed on mature GFP/MAP2 positive neurons which showed significantly larger diameter of T21 dendrites at DIV100 (Fig. 3g). Although the T21 dendrites were shorter (Fig. 3h), with fewer branches (Fig. 3i) compared to the D21 dendrites, the differences were not significant. Importantly, the same morphological features of cortical neurons were observed in both, infant and adult age- and sex- matched human brain (Fig. 3j, Supplementary Table 1). T21 neurons showed significantly larger diameter of MAP2 positive dendrites (Fig. 3k) which were at the same time shorter (Fig. 3l) with significantly fewer branches (Fig. 3m) compared to euploid controls. Our data showed the same pattern in 2D neurons and in the human brain, while in mature hCSAs, neurons exhibited the same trend, although with lower statistical significance. This result can be explained by differences in cell density and tissue organisation. Neurons on coverslips were allowed the space to more evenly distribute across the surface, whereas in the human brain, cortical neurons were organised in distinct layers. In contrast, neurons in hCSAs were densely packed in 3D structures, but they lack clear layering and organisation, which may be a disadvantage for dendrite analyses. Taken together, the abnormal neuronal phenotype caused by T21 was reflected in the following measurements: (1) T21 neurons had up to a 1.4-fold larger diameter (Fig. 3b, g, k), (2) T21 neurons were up to 1.36-fold shorter (Fig. 3c, h, l) and (3) T21 neurons contained up to 1.33-fold fewer branches (Fig. 3d, i, m) compared to D21 neurons in both, the isogenic system and in human brain. Next, we analysed the branching points, terminal points and the Sholl diagram in both 2D neurons (Supplementary Fig. 3a-c) and hCSAs (Supplementary Fig. 3d-f), as well as in the human brain (Supplementary Fig. 3g-i) and observed a similar pattern, although the effects were less significant. Despite the fold differences being even higher for these measurements, reaching up to 1.5-fold lower values in T21 neurons, the statistical significance was reduced due to the broader distribution of neurons. Finally, the same approach was used to analyse isogenic axons in 2D. Interestingly, T21 SMI-312 positive axons showed a significantly larger diameter compared to the D21 axons, reaching up to 1.15-fold larger values in T21 neurons (Supplementary Fig. 3j-k).

**Fig. 3.**
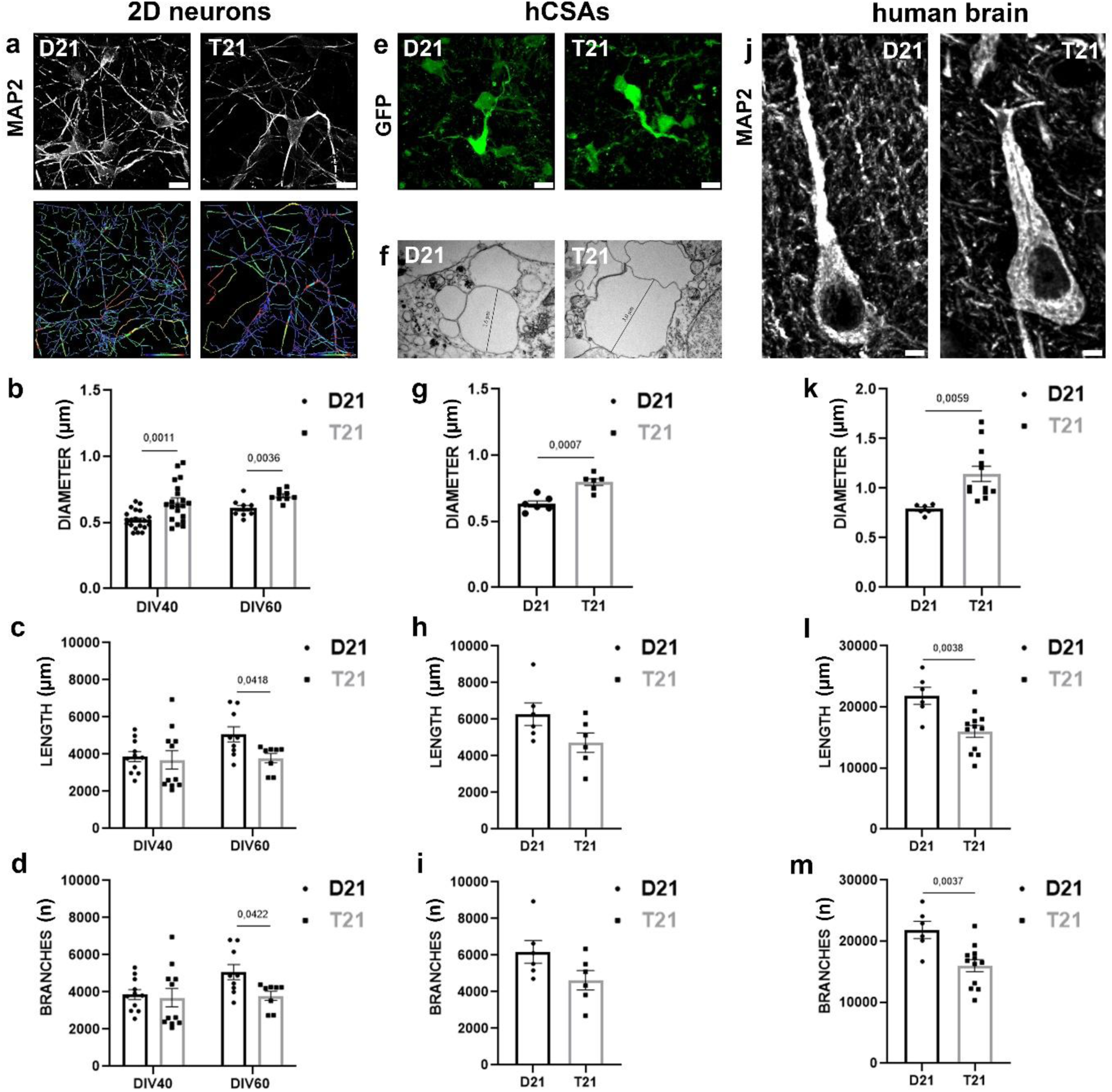
**T21 neurons showed abnormal neuronal morphology**. **(a-d)** 2D neurons: **(a)** Representative confocal images of isogenic 2D neurons and the same-Imaris 9.9.1 processed images. Different colour of neurites were Imaris software coded and represents diameter, length and branches. Scale bar: 10 µm. **(b)** T21 dendrites showed significantly larger diameter, **(c)** T21 dendrites were significantly shorter with **(d)** significantly fewer branches compared to the D21 dendrites. (**e-i)** hCSAs: **(e)** Representative confocal images of isogenic hCSAs neurons. Scale bar: 10 µm. **(f)** Representative TEM images of isogenic dendrites, magnification 20,000x. The representative diameter of the T21 dendrite measured 3.0 µm, whereas the D21 dendrite measured 2.6 µm. **(g)** T21 dendrites showed significantly larger diameter, **(h)** T21 dendrites were shorter with **(i)** fewer branches compared to the D21 dendrites. **(j-m)** human brain: **(j)** Representative confocal images of human cortical neurons. **(k)** T21 dendrites showed significantly larger diameter, **(l)** T21 dendrites were significantly shorter with **(m)** significantly fewer branches compared to the D21 dendrites. Dendrite diameter and length are presented in µm per field of view (y-axis), while branches are presented as the number (n) of branches per field of view. The x-axis represents DIV40 and DIV60 for 2D neurons **(b–d)**, whereas for assembloids **(g–i)** and human brain samples **(k–m)**, it represents genotype. Graphs represent means ± SEM.

Additionally, isogenic hCSs and hStrSs at DIV50, as well as hCSAs at DIV100, were analysed using scanning electron microscopy (SEM). Our data revealed abnormal neuronal morphologies caused by T21 in hCSs, hStrSs, and hCSAs throughout differentiation. T21 neurons exhibited shorter neurites with a larger diameter, less branches and less dendritic spines, whereas D21 neurons were evenly distributed and displayed a robust neuronal network composed of thin, long, highly branched neurites with dense dendritic spines. Overall, our SEM data demonstrated a more compact and dense tissue organisation in D21 spheroids/assembloids compared to T21 (Fig. 4). Guided by the morphological data, we investigated whether our isogenic system exhibits detectable differences in mitochondria and synapses.

**Fig. 4.**
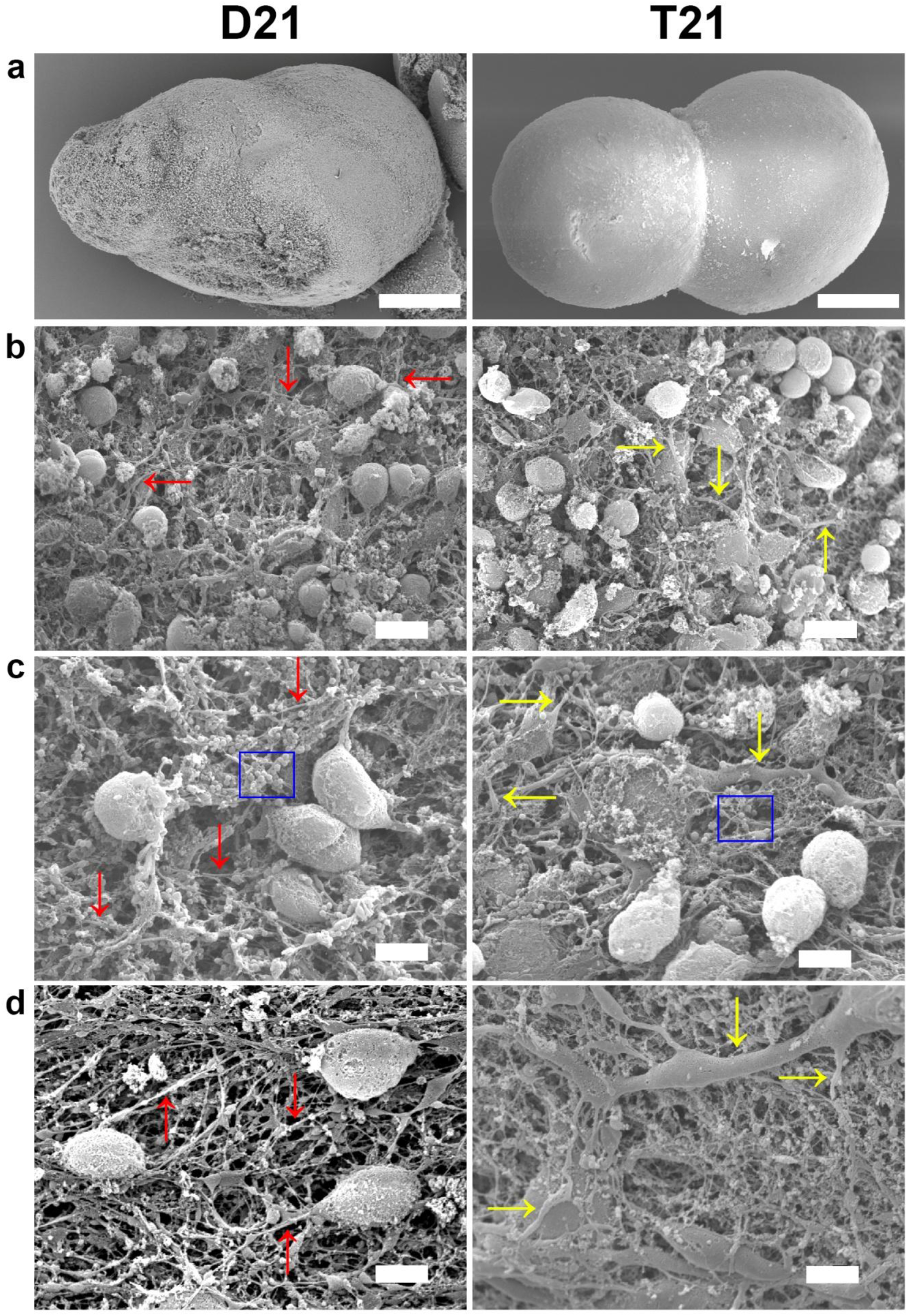
**High resolution imaging of isogenic neurons**. Representative SEM images of isogenic hCSAs at DIV100, captured at various magnifications, revealed abnormal neuronal morphology represented by shorter neurites with larger diameter, fewer branches and lower density of synaptic spines in T21 hCSAs. Legend: D21 neurites (red arrows), T21 neurites (yellow arrows) and dendritic spines (blue square). Scale bar: **(a)** 100 µm, **(b)** 4 µm and **(c-d)** 2 µm.

### T21 cells showed abnormal mitochondrial morphology

As neurite branching and termini formation are linked to the biology and cellular distribution of mitochondria [20–22], we next, analysed mitochondrial morphology in our isogenic system. Our data showed that 80.9% T21 NSCs and 92.8% D21 NSCs contain MitoTracker positive mitochondria (Fig. 5a-b), indicating a significantly lower proportion of functionally active mitochondria in T21 cells. Interestingly, significantly higher expression of AIF, a marker of the inter-membrane space, was observed in T21 cells, normalised by ß-actin (Fig. 5c-d) and Nestin (Fig. 5e-f). Similarly, significantly higher expression of TOMM20, a marker of the outer mitochondrial membrane (Fig. 5g-h) was measured in T21 cells. Pairwise Pearson’s coefficient of colocalization showed a high level of colocalization (0.7) between AIF and TOMM20 (Fig. 5i). A significantly higher level of AIF relative to MAP2 was found between T21 and D21 neurons in 2D (Fig. 5j-k) which also appears to be observed in hCSAs (Fig. 5l). Interestingly, in T21 neurons most mitochondria were aggregated within the perikaryon, while in D21 neurons they were evenly distributed throughout the cell. Finally, abnormal mitochondrial morphology was confirmed by TEM on isogenic hCSAs at DIV100 (Fig. 5m). Overall, our data showed a significantly higher number of functionally active, MitoTracker positive mitochondria in D21 cells. At the same time, our data showed mitochondrial aggregation and clumping in all T21 cells what causes significantly higher expression of AIF and TOMM20 throughout *in vitro* differentiation. This likely results from disrupted mitochondrial fusion and fission processes, leading to mitochondrial fragmentation, a feature characteristic of many neurodegenerative disorders, including DS. Our study suggests the onset of these abnormalities to be in neurodevelopmental period, potentially pre-dating the obvious patho-histological signs of neurodegeneration in DS. These findings encouraged the further study of synaptogenesis in our system as mitochondria play an essential role in synaptic formation and function.

**Fig. 5.**
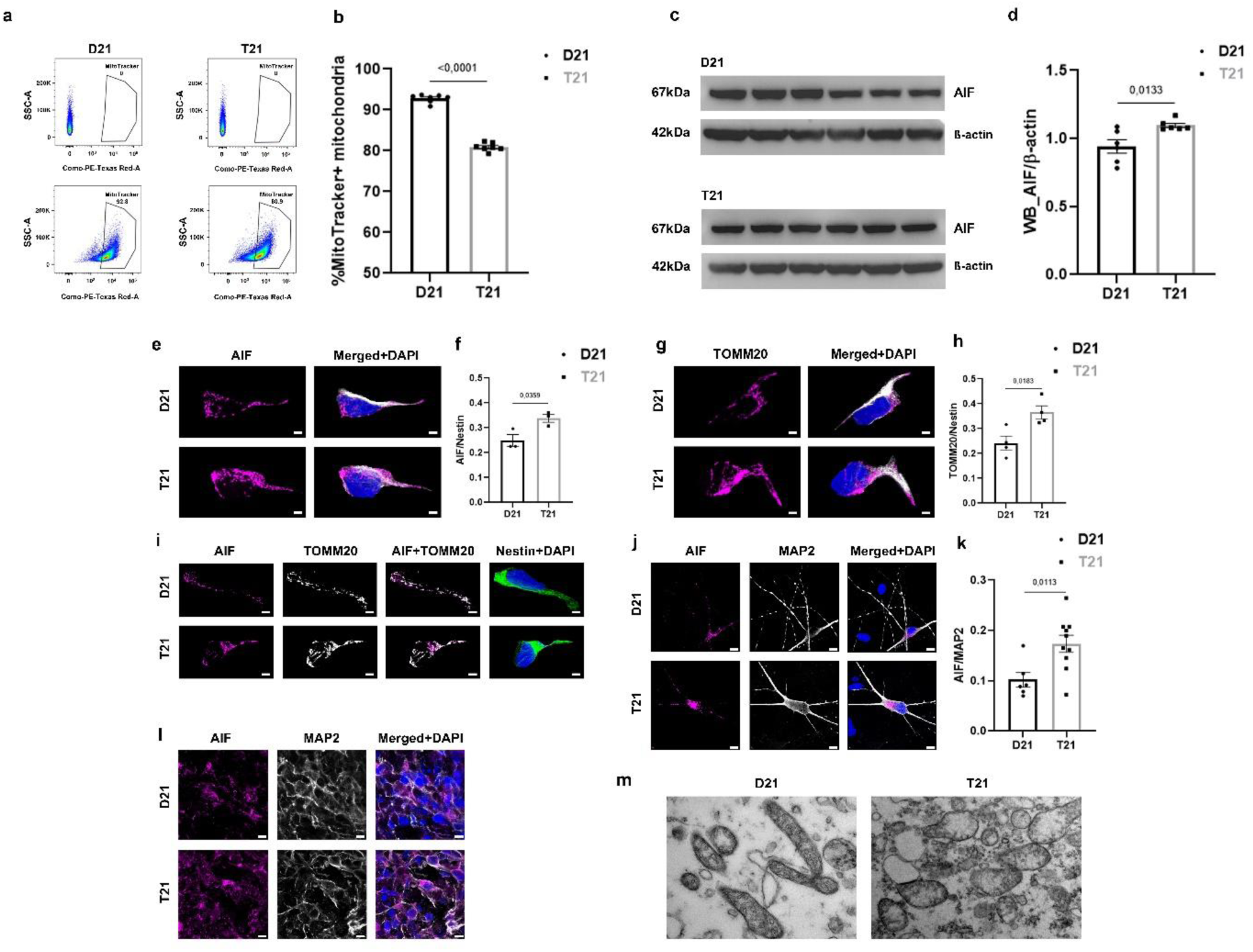
Isogenic T21 cells showed abnormal mitochondrial morphology. (a-k) 2D cultures: **(a)** Representative plots of unlabelled and MitoTracker (functionally active mitochondria) labelled isogenic NSCs showed 80.9% labelled mitochondria in T21 NSCs and 92.8% labelled mitochondria in D21 NSCs analysed by flow cytometry. The y-axis (SSC-A) represents signal area on side scatter (SSC) detector, while x-axis represents fluorescent intensity of MitoTracker signal area. **(b)** T21 NSCs showed significantly lower percentage of MitoTracker positive mitochondria. The y-axis represents percentage of MitoTracker positive mitochondria, while x-axis represents genotype. **(c)** Representative blots of AIF/ß-actin of isogenic NSCs. **(d)** WB analysis of AIF/ß-actin showed significantly higher expression in T21 NSCs. The y-axis represents expression of AIF normalised by β-actin, while x-axis represents genotype. **(e)** Representative confocal images of NSCs stained with AIF (magenta), Nestin (white) and DAPI (blue). Scale bar: 2 µm. **(f)** T21 NSCs showed significantly higher AIF expression. The y-axis represents expression of AIF normalised by Nestin, while x-axis represents genotype. **(g)** Representative confocal images of NSCs stained with TOMM20 (magenta), Nestin (white) and DAPI (blue). Scale bar: 2 µm. **(h)** T21 NSCs showed significantly higher TOMM20 expression. The y-axis represents expression of TOMM20 normalised by Nestin, while x-axis represents genotype. **(i)** Representative confocal images of colocalization between AIF (magenta) and TOMM20 (white) counterstained with Nestin (green) and DAPI (blue) showed high Pairwise Person’s coefficient of colocalization (0.7). **(j)** Representative confocal images of AIF positive mitochondria (magenta) in MAP2 (white) positive neurons at DIV60. Scale bar: 10 µm. **(k)** T21 neurons showed significantly higher AIF expression. The y-axis represents expression of AIF normalised by MAP2, while x-axis represents genotype. **(l-m)** hCSAs: **(l)** Representative confocal images of mature hCSAs at DIV100 stained with AIF (magenta), MAP2 (white) and DAPI (blue). Scale bar: 10 µm. **(m)** Representative TEM images of isogenic mitochondria in hCSAs showed abnormal mitochondrial morphology in T21 cells at DIV100, magnification 50,000x. Graphs represent means ± SEM.

### T21 neurons showed abnormal synaptic morphology throughout *in vitro* differentiation

After analysing neuronal morphology, we aimed to compare T21 and D21 synaptogenesis throughout *in vitro* differentiation (Fig. 6, Supplementary Figs. 4-5). For the purpose of this study, we analysed the presynaptic marker (SYN-1) and two postsynaptic markers (PSD95 and GEPH). In T21 cells, synaptic puncta formed large, aberrant clumps, whereas in D21 cells both, presynaptic and postsynaptic puncta were evenly distributed across neurites (Fig. 6a). The expression of all three synaptic genes was significantly lower in T21 neurons throughout differentiation (Fig. 6b). Interestingly, the levels of PSD95 and SYN-1 increased dramatically from DIV40 to DIV60 in both T21 and D21 neurons, while the level of GEPH remained unchanged (Fig. 6c, Supplementary Figs. 4-5). These results were confirmed in total cell lysates at the same time points (Fig. 6d-e, Supplementary Fig. 4). Our data showed significantly less PSD95, GEPH and SYN-1 puncta in T21 dendrites (Fig. 6f-g, Supplementary Figs. 4-5) and in T21 axons (Fig. 6f, h, Supplementary Figs. 4-5) compared to D21 dendrites and axons at DIV40 and DIV60.

**Fig. 6.**
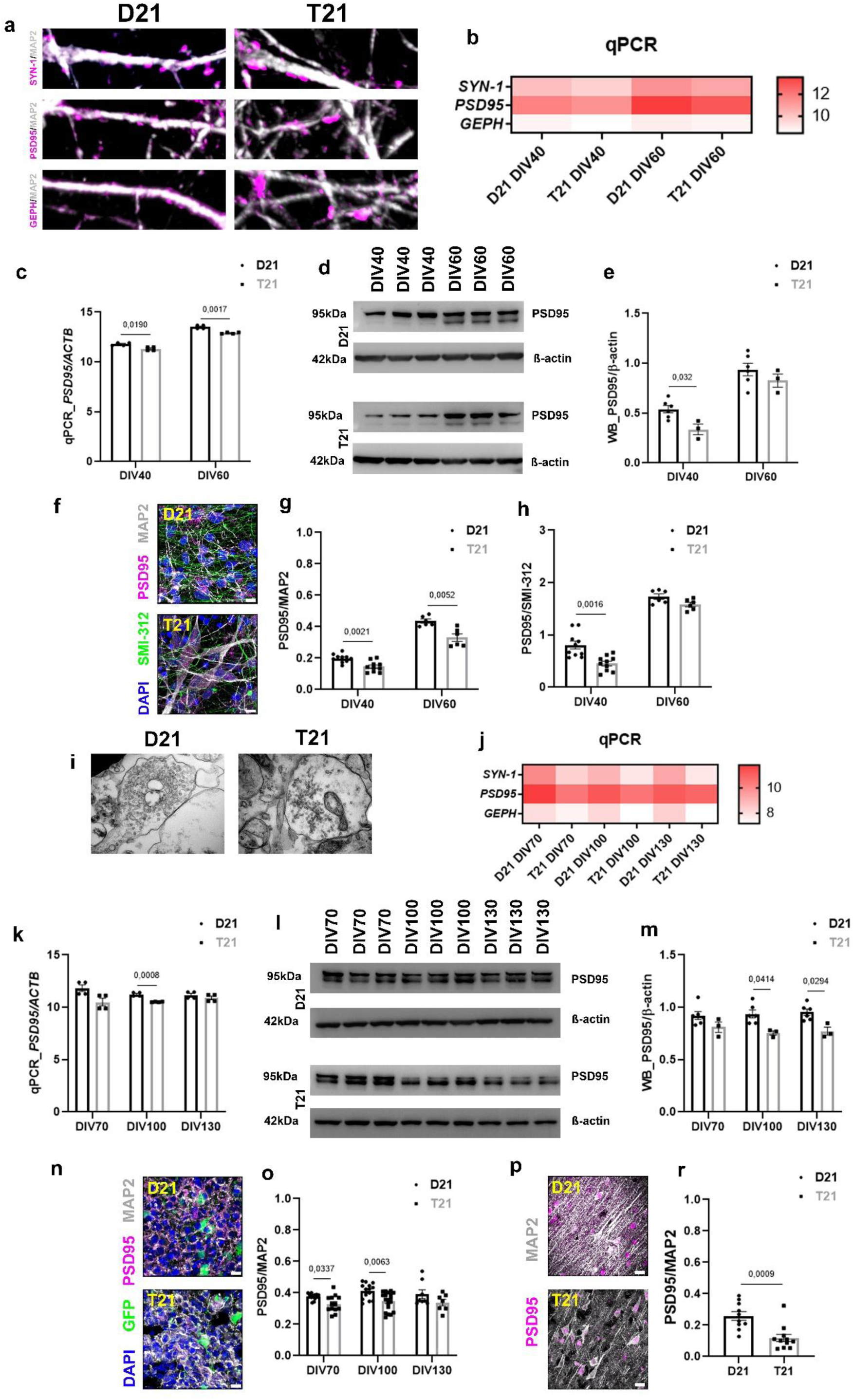
**Isogenic T21 neurons showed reduced synaptogenesis and abnormal synaptic morphology**. **(a-h)** 2D neurons: **(a)** Representative confocal images of SYN-1, PSD95 and GEPH positive synaptic puncta (magenta) in mature MAP2 (white) positive isogenic dendrites at DIV60. T21 synaptic puncta formed aberrant clumps while the D21 synaptic puncta were evenly distributed across dendrites, magnification 60x. **(b)** Gene expression of all three synaptic markers throughout *in vitro* differentiation. **(c)** Analysis of *PSD95/ACTB* throughout *in vitro* differentiation showed the lower gene expression of *PSD95* in T21 neurons. The y-axis represents gene expression of *PSD95* normalised by *ACTB*, while x-axis represents DIV40 and DIV60. **(d)** Representative blots of PSD95/ß-actin throughout *in vitro* differentiation. **(e)** WB analysis of PSD95/ß-actin showed significantly lower expression in T21 neurons. The y-axis represents expression of PSD95 normalised by β-actin, while x-axis represents DIV40 and DIV60. **(f)** Representative confocal images of mature isogenic 2D neurons stained with SMI-312 (green), PSD95 (magenta), MAP2 (white) and DAPI (blue) at DIV60. Scale bar: 10 µm. **(g)** T21 neurons showed significantly less PSD95 positive synaptic puncta in MAP2 positive dendrites and **(h)** SMI-312 positive axons throughout *in vitro* differentiation compared to D21 neurons. The y-axis represents expression of PSD95 normalised by MAP2 **(g)** and SMI-312 **(h)**, while x-axis represents DIV40 and DIV60. **(i-o)** hCSAs: **(i)** Representative TEM images of isogenic hCSAs showed distribution of synaptic puncta at DIV100, magnification 50,000x. **(j)** Gene expression of all three synaptic markers throughout hCSAs differentiation. **(k)** Analysis of *PSD95/ACTB* showed significantly lower gene expression in T21 hCSAs compared to D21 hCSAs throughout *in vitro* differentiation. The y-axis represents gene expression of *PSD95* normalised by *ACTB*, while x-axis represents DIV70, DIV100 and DIV130. **(l)** Representative blots of PSD95/ß-actin throughout *in vitro* differentiation. **(m)** WB analysis of PSD95/ß-actin showed significantly lower expression in T21 hCSAs. The y-axis represents expression of PSD95 normalised by β-actin, while x-axis represents DIV70, DIV100 and DIV130. **(n)** Representative confocal images of mature isogenic hCSAs with GFP positive striatal neurons (green), stained with PSD95 (magenta), MAP2 (white) and DAPI (blue) at DIV100. Scale bar: 10 µm. **(o)** T21 neurons showed significantly less PSD95 positive synaptic puncta in MAP2 positive dendrites throughout *in vitro* differentiation compared to D21 neurons. The y-axis represents expression of PSD95 normalised by MAP2, while x-axis represents DIV70, DIV100 and DIV130. **(p-r)** human brain: **(p)** Representative confocal images of cortical neurons from human brains stained with PSD95 (magenta) and MAP2 (white). Scale bar: 10 µm. **(r)** T21 neurons showed significantly less PSD95 positive synaptic puncta in MAP2 positive cortical dendrites compared to D21 cortical neurons. The y-axis represents expression of PSD95 normalised by MAP2, while x-axis represents genotype. Graphs represent means ± SEM.

Additionally, the same analysis was performed on hCSAs at DIV70, DIV100 and DIV130. In order to show the distribution of synaptic puncta across the neurites, TEM analysis was performed at DIV100. Similar to 2D neurons, T21 hCSAs neurites displayed abnormal clumps and more sparsely distributed puncta, whereas D21 hCSAs exhibited evenly distributed and dense synaptic puncta compared to the isogenic control (Fig. 6i). The expression of all three synaptic marker genes was significantly lower in T21 hCSAs throughout differentiation (Fig. 6j). Our data showed significantly lower expression of PSD95 in T21 hCSAs analysed by qPCR (Fig. 6k), WB (Fig. 6l-m) and immunofluorescence (Fig. 6n-o). Moreover, the same results were obtained with another postsynaptic marker GEPH (Supplementary Fig. 4) as well as presynaptic marker SYN-1 (Supplementary Fig. 5). Put together, our data suggest that synaptogenesis was reduced in T21 neurons compared to D21 neurons at the same stage of differentiation in both 2D neurons and hCSAs. Importantly, the reproducible reduction in PSD95 expression observed in isogenic T21 neurons throughout *in vitro* differentiation was confirmed in the human brain (Fig. 6p-r, Supplementary Table 1).

### T21 spheroids were smaller compared to D21 spheroids

During the culturing of isogenic spheroids/assembloids, we observed distinct size differences between T21 and D21 spheroids/assembloids. To analyse the circumference and area of spheroids using Lusca software throughout the first 40 days of *in vitro* differentiation we analysed both isogenic hCSs (Fig. 7a-b) and isogenic hStrSs (Fig. 7c-d) prior to the fusion into assembloids. At each time point, five to seven images were captured per clone/spheroid, resulting in a total analysis of over 200 spheroids. Although all spheroids were initially seeded with the same number of cells (10,000/well), significant size differences were observed. As expected, there were no differences in circumference and area for both T21 and D21 spheroids at DIV1 (Supplementary Fig. 6). However, from DIV5 to DIV40, highly significant differences emerged between the clones. In this study, we aimed to compare both parameters, circumference and area, between T21 and D21 spheroids. First, we compared the circumference (Fig. 7e) and area (Fig. 7f) of T21 and D21 hCSs. Our data showed significant differences as early as DIV1, with these differences increasing throughout *in vitro* differentiation. These disparities can likely be attributed to the superior spheroid-forming capacity and higher survival of D21 cells. Specifically, throughout differentiation, the circumference of T21 hCSs was 10-15% smaller than circumference of D21 hCSs, depending on the time point (Fig. 7e), while the area of T21 hCSs was 20-30% smaller than the area of D21 hCSs (Fig. 7f). Interestingly, a similar trend was observed in hStrSs, although the differences were not highly significant across all time points (Fig. 7g-h). Nonetheless, the circumference of T21 hStrSs was consistently 13-18% smaller than the circumference of D21 hStrSs throughout *in vitro* differentiation (Fig. 7g), while the area of T21 was 20-27% smaller (Fig. 7h) compared to D21. The same pattern was observed after fusion into the hCSAs until DIV130 (Fig. 2a, Fig. 4a, Fig. 7i-j), but investigated parameters were not quantified.

**Fig. 7.**
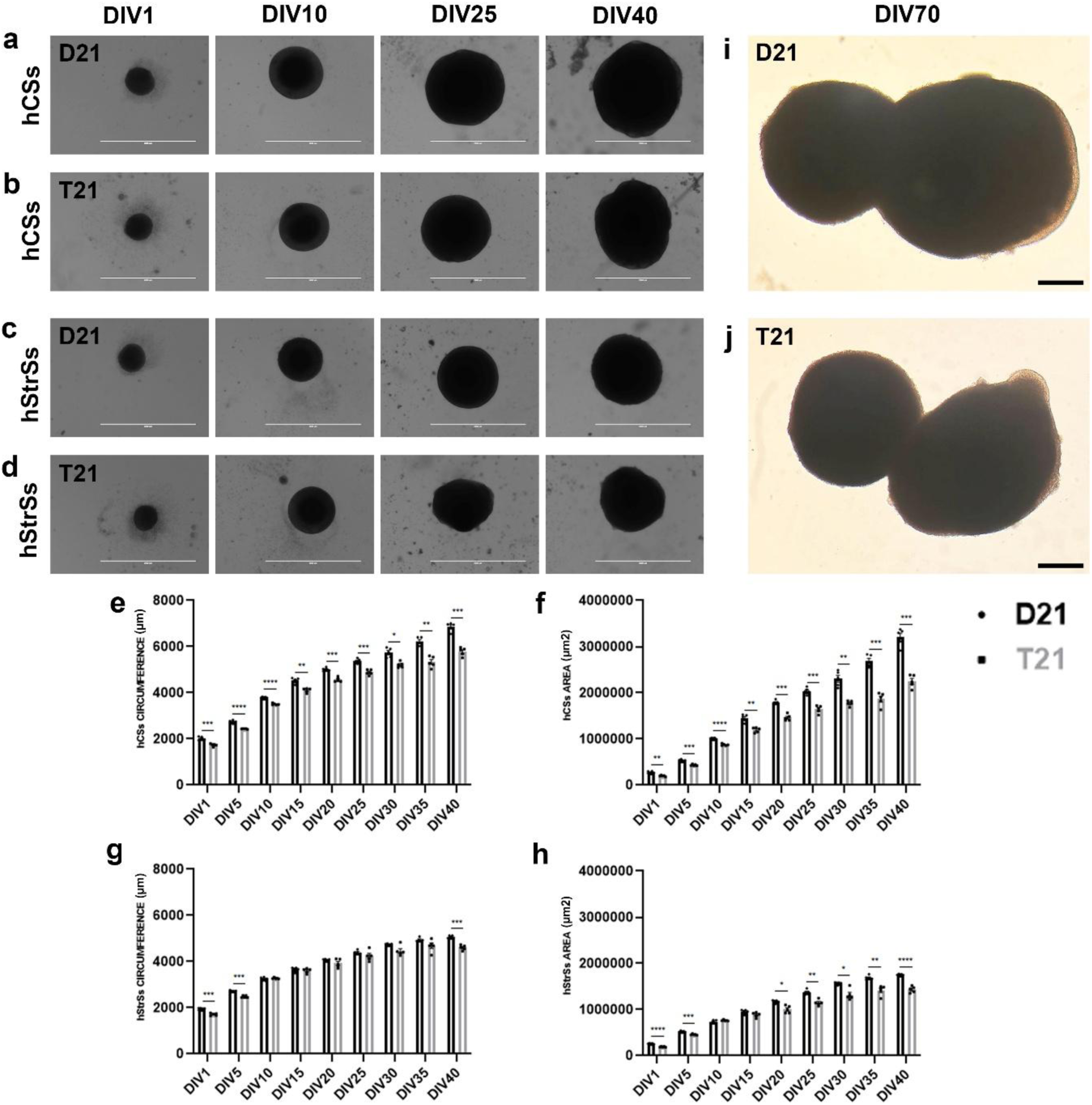
**T21 spheroids were significantly smaller compared to D21 spheroids**. Representative images of **(a)** D21 hCSs, **(b)** T21 hCSs, **(c)** D21 hStrSs and **(d)** T21 hStrSs at DIV1, DIV10, DIV25 and DIV40. Scale bar: 2000 µm. **(e)** Comparison of the T21 and D21 hCSs circumference showed significantly lower values in T21 spheroids throughout *in vitro* differentiation. **(f)** Comparison of the T21 and D21 hCSs area showed significantly lower values in T21 spheroids throughout *in vitro* differentiation. **(g)** Comparison of the T21 and D21 hStrSs circumference showed lower but less significant values in T21 spheroids throughout *in vitro* differentiation. **(h)** Comparison of the T21 and D21 hStrSs area showed lower but less significant values in T21 spheroids throughout *in vitro* differentiation. **(i)** Representative images of D21 and **(j)** T21 hCSAs at DIV70. Scale bar: 500 µm. Circumference **(e, g)** is presented in µm (y-axis), while area **(f, h)** is presented in µm^2^ (y-axis). The x-axis represents DIV1-40 **(e-h)**. Graphs represent means ± SEM.

Additionally, when we analysed the same parameters in T21 and D21 spheroids separately, we found that the circumference of both T21 and D21 hCSs increased 3.4-fold throughout *in vitro* differentiation, whereas the circumference of T21 hStrSs increased 2.7-fold and that of D21 hStrSs increased 2.5-fold (Supplementary Fig. 6a-b). In contrast, the area of T21 hCSs increased 11.3-fold and that of D21 hCSs increased 11.6-fold, while the area of T21 hStrSs increased 7.5-fold and that of D21 hStrSs increased 8.75-fold throughout *in vitro* differentiation (Supplementary Fig. 6c-d).

Overall, our results indicate that the spheroids reflect patterns of cell differentiation consistent with human brain development, wherein the cortex is significantly larger than the striatum and DS brains are smaller compared to euploid controls.

### T21 hCSAs contain more dead cells and less proliferating cells

Next, we investigated whether the cell proliferation could explain morphological readouts. We analysed the number of live/dead cells at DIV1 and DIV5, as well as quantified expression of SOX2, Nestin, Ki67, CCaspase-3 and performed a TUNEL assay in both spheroids and hCSAs throughout *in vitro* differentiation. Given the significant differences in spheroid morphology observed at DIV1 and DIV5 (Fig. 7e-h) between T21 and D21 spheroids, we analysed cell viability using flow cytometry. Our data showed that T21 spheroids contained 4.3% fewer live cells at DIV1 and 21.95% fewer live cells at DIV5 than D21 spheroids (Fig. 8a). As expected, SOX2 expression decreased throughout *in vitro* differentiation in D21 spheroids, but the expression took longer to decrease in T21 spheroids (Supplementary Fig. 7a-c). Comparison of T21 and D21 spheroids, showed a similar SOX2 expression at DIV30 in both isogenic hCSs and hStrSs, with a 1.6-fold higher expression in hCSs. Interestingly, differentiation resulted in significantly more SOX2 positive cells in both T21 spheroids, hCSs and hStrSs (Fig. 8b-c, Supplementary Fig. 7a-c), while Nestin showed the same level of expression at DIV70 (Supplementary Fig. 7d-e). Then we analysed SOX2 expression in hCSAs throughout differentiation; our data showed similar level of expression in T21 hCSAs, while in D21 samples we showed significant decrease throughout differentiation (Fig. 8d-e).

**Fig. 8.**
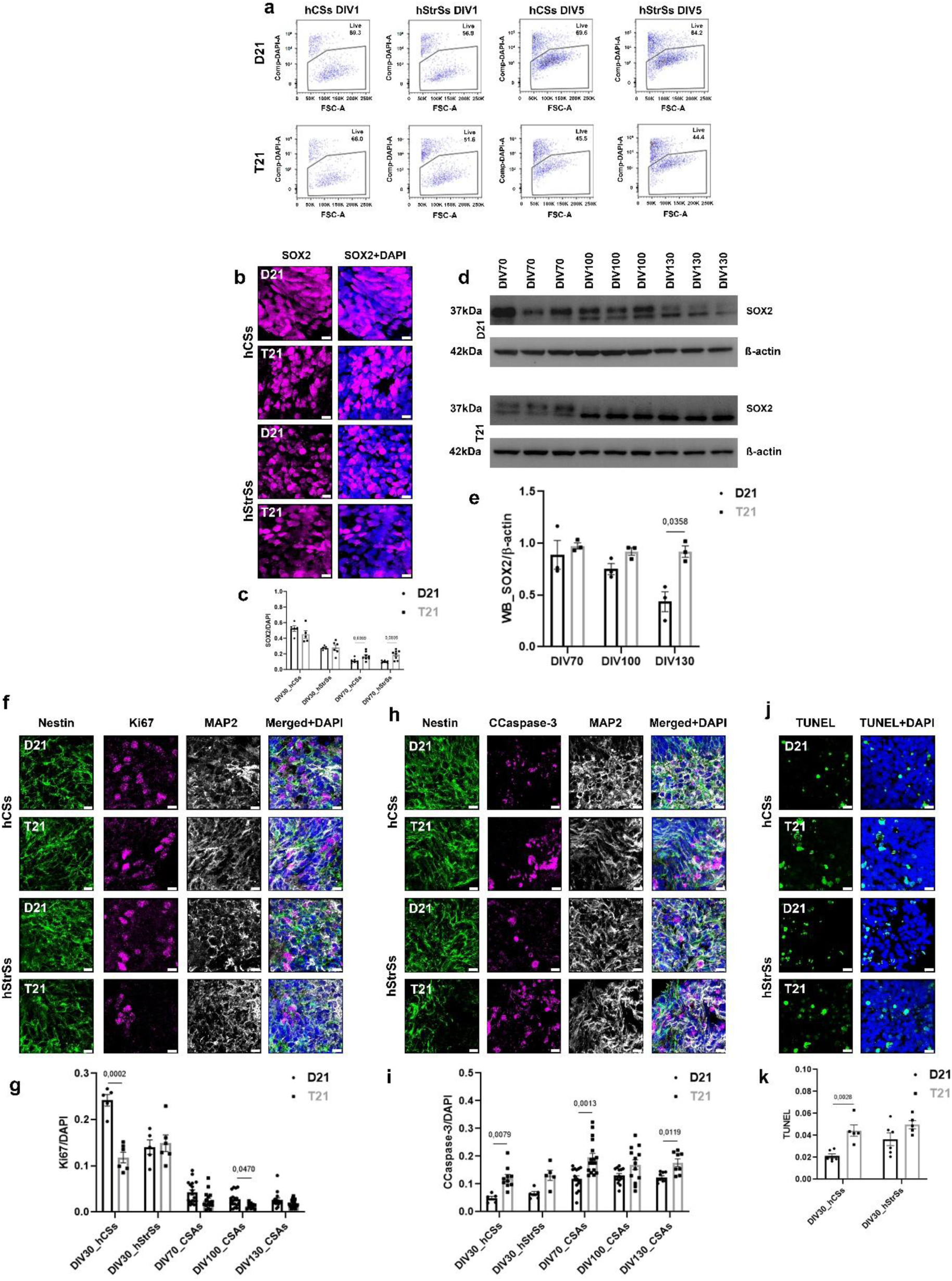
**Proliferation and cell death throughout *in vitro* differentiation**. **(a)** Analysis of live/dead cells by flow cytometry using DAPI: representative plots of hCSs and hStrSs showed 4.3% fewer live cells in T21 spheroids at DIV1 and 21.95% fewer live cells in T21 spheroids at DIV5. The y-axis represents fluorescent intensity of DAPI, while x-axis (FSC-A) represents area of forward scatter (FSC) detector. **(b)** Representative confocal images of isogenic hCSs and hStrSs stained with SOX2 (magenta) and DAPI (blue) at DIV30. Scale bar: 10 µm. **(c)** Isogenic spheroids showed similar expression of SOX2 at DIV30, while at DIV70 T21 spheroids showed significantly more SOX2 positive cells. The y-axis represents expression of SOX2 normalised by DAPI, while x-axis represents DIV30 and DIV70. **(d)** Representative blots of SOX2/ß-actin throughout *in vitro* differentiation. **(e)** WB analysis of SOX2/ß-actin showed higher expression of SOX2 in T21 hCSAs throughout *in vitro* differentiation. The y-axis represents expression of SOX2 normalised by β-actin, while x-axis represents DIV70, DIV100 and DIV130. **(f)** Representative confocal images of isogenic hCSs and hStrSs stained with Nestin (green), Ki67 (magenta), MAP2 (white) and DAPI (blue) at DIV30. Scale bar: 10 µm. **(g)** T21 cells showed significantly fewer proliferating cells labelled with Ki67 throughout *in vitro* differentiation. The y-axis represents expression of Ki67 normalised by DAPI, while x-axis represents DIV30, DIV70, DIV100 and DIV130. **(h)** Representative confocal images of isogenic spheroids stained with Nestin (green), CCaspase-3 (magenta), MAP2 (white) and DAPI (blue) at DIV30. Scale bar: 10 µm. **(i)** T21 spheroids and hCSAs showed significantly more dead cells labelled with CCaspase-3 throughout *in vitro* differentiation. The y-axis represents expression of CCaspase-3 normalised by DAPI, while x-axis represents DIV30, DIV70, DIV100 and DIV130. **(j)** Representative confocal images of isogenic spheroids stained with TUNEL (green) and DAPI (blue) at DIV30. Scale bar: 10 µm. **(k)** T21 spheroids showed significantly more dead cells at DIV30. The y-axis represents expression of TUNEL positive cells normalised by DAPI, while x-axis represents DIV30. Graphs represent means ± SEM.

Furthermore, analysis of proliferating cells labelled with Ki67 revealed significant differences across all investigated time points, with lower expression in T21. Reduction in Ki67 throughout the differentiation timeline would be expected, as neurons traject towards a post-mitotic stage. However, the T21 hCSAs still showed significantly less Ki67 positive cells, and this could be attributed to the causative effect by trisomy 21 (Fig. 8f-g, Supplementary Fig. 6f-h).

Cell death was assessed using CCaspase-3 and TUNEL assay. In contrast to Ki67, expression of CCaspase-3 was significantly higher in T21 at all time points. Interestingly, the lowest expression was observed in spheroids at DIV30, while in hCSAs, expression remained stable from DIV70 to DIV130 (Fig. 8h-i, Supplementary Fig. 6i-k). Additional confirmation of cell death was provided by a TUNEL assay in spheroids at DIV30 (Fig. 8j-k). When the ratio of proliferation to cell death was calculated, it became evident that in T21 hCSAs, up to ten times more cells died than proliferated throughout *in vitro* differentiation. Contrary to this, in D21 hCSAs, twice as many cells proliferated than died.

Altogether, based on our data, we conclude that abnormal neuronal morphology is caused by T21, and this, along with reduced proliferation and increased cell death, leads to significantly smaller T21 spheroids - mirroring observations in the developing human brain of DS.

## DISCUSSION

Our study identifies fundamental effects of T21 on neuronal morphology throughout *in vitro* differentiation, reproducing the effects visible in the human brain. Several studies have reported different morphological features linked to T21 in ESCs [29], primary neurons [23, 24], interneurons [35] and mouse neurons [36–38]. However, very few studies have examined morphological differences in isogenic iPSC systems, that can causatively be attributed to T21, allowing further modelling *in vitro*. Ability to faithfully reproduce these neurodevelopmental defects in 2D and 3D systems derived from iPSCs, opens possibilities of assigning the specific phenotypes to specific genes which are overdosed in trisomy 21. The limited existing work has shown that DS GABA neurons are smaller and have fewer neuronal processes [27], but their detailed morphology has not yet been described. Using isogenic iPSCs [3, 30] we showed specific neuronal morphology caused by T21. The main morphological feature of our isogenic system in 2D was that T21 neurons formed aggregates of cell bodies, whereas D21 neurons were evenly distributed across the surface, forming a strong neuronal network. However, in addition to these aggregates of cell bodies in T21 cultures, some neurons were distributed in between them and exhibited the following features: (1) T21 dendrites showed significantly larger diameter, (2) T21 dendrites were significantly shorter, (3) T21 dendrites showed significantly fewer branches, (4) T21 dendrites showed significantly fewer branching points, (5) T21 dendrites showed significantly fewer terminal points and (6) T21 dendrites showed significantly smaller field size through the Sholl analysis. Additionally, we wanted to see if the same phenotype was reproducible in 3D structures such as hCSAs [31, 32]. In isogenic hCSAs at DIV100, as well as in foetal and postnatal human brain samples we observed the same pattern. Importantly, here we show the same morphological features in both the 2D and 3D isogenic system as well as in the human brain. Although we focused on neurons and their morphology, in our conditions we observed significantly more T21 astrocytes in both the 2D and 3D systems; a finding that has been previously reported in cerebral organoids [39], the human brain [40] and rodent models of DS [41, 42]. These findings suggest a pivotal role for astrocytes throughout neuronal *in vitro* differentiation, as also observed in the human brain [40, 43], particularly with the specific glial shift seen in DS.

Neurofilaments are proteins of the nervous system that form part of the neuronal cytoskeleton and play essential roles in regulating microtubule and organelle dynamics, nerve conduction, and neurotransmission [44, 45]. In our study, we focused on neuronal morphology and its close relationship with mitochondria and synaptogenesis in our isogenic model. The results showed abnormal mitochondrial morphology particularly, the formation of clumps and aggregations, consistent with findings from previous studies on human cells [46–48] and mouse models of DS [49, 50]. Since mitochondria are involved in synaptic transport, neurotransmitter formation and provide energy for these processes [51], we analysed synaptogenesis in both 2D and 3D systems. Our study revealed a significant reduction in both presynaptic and postsynaptic puncta throughout differentiation of 2D neurons and hCSAs. Moreover, our recent study demonstrated a reduced electrophysiological activity of excitatory T21 neurons cultured in 2D [17], consistent with a previous study which showed reduced synaptic density in DS brain [52]. Some of the other studies showed reduced neuronal activity of hiPSCs derived T21 neurons after transplantation into the mouse brain [53, 54] as well as altered neurophysiology of hippocampal neurons in a mouse model of DS [55]. Finally, in addition to abnormal mitochondrial and synaptic morphology, enlarged endosomes in T21 neurons [56], cerebral organoids [39] and mouse basal forebrain neurons [38] were associated with impaired neuronal trafficking.

Abnormal neuronal morphology, along with abnormal mitochondria and reduced synaptogenesis caused by T21 in our system, was accompanied by decreased proliferation and increased cell death. The results of our study showed significantly smaller T21 spheroids compared to D21 spheroids throughout *in vitro* differentiation. Similar observations have previously been reported in T21 cerebral organoids [57], as well as in partial T21 cerebral organoids and their corrected controls [28, 30]. Moreover, T21 cells showed significant migration deficits of neural crest [58]. This finding corresponds to the development of the human DS brain during gestation and postnatal life, which is characterised by an overall smaller brain volume, reduced grey and white matter, diminished gyrification, and other brain abnormalities [59–62]. Here we showed that T21 hCSAs have fewer dendritic spines at DIV100 using scanning electron microscopy, which is consistent with a previous study of the adult human brain stained with classical Golgi staining [63]. In our previous study on cerebral organoids, we demonstrated the correction of “gyrification” following CRISPR-mediated correction of the *DYRK1A* gene in partial T21 cells [28]. Moreover, at the cellular level, we demonstrate that T21 spheroids contained significantly more CCaspase-3 positive (dead cells) cells compared to D21 spheroids throughout differentiation. At the same time, D21 spheroids exhibited significantly more Ki67 positive (proliferating cells) cells compared to T21 spheroids. These findings correspond to a previous published study on the human foetal hippocampus (17-21 weeks of gestation) [40, 57]. Besides the potential role of the triplication of *DYRK1A*, it is tempting to speculate which other HSA21 genes might be involved. A result diametrically opposite to ours (i.e. a severe acceleration of neuronal differentiation, resulting in the premature loss of SOX2+ progenitors, and accelerated accumulation of post-mitotic neuronal differentiation markers, such as DCX) was recently observed in hiPSCs in which *APP* gene was knocked out using CRISPR/Cas9 (APP^null^) [64]. It is tempting to speculate that trisomy of APP gene (i.e. DS) might produce the opposite effects. However, other overdosed genes must certainly be involved, as people born with the duplication of *APP* gene, despite having an obligatory early-onset AD, are not known to suffer from obvious clinical neurodevelopmental defects.

The main limitation of our study is the use of a single isogenic system, even though the study was performed using three D21 and three T21 iPSC clones [3]. Additionally, one D21 and one T21 line contained fully integrated GFP [30]. In total, eight isogenic iPSC lines were used and differentiated into 2D neurons and hCSAs. The potential bias of the results based on an isogenic iPSC system from a single individual is overcome by the reproduction of phenotypes in human DS brain samples, using the same assessment methods. Although we were limited by the availability of age- and gender-matched human brain samples, in total, we used tissue from fifteen individuals from the Zagreb Neuroembryological Collection of the Croatian Institute for Brain Research (CIBR) [33]. Future work should involve single-cell analysis of isogenic spheroids and hCSAs, followed by analysis of cell-to-cell interactions, neuronal circuit formation, and maturation in the DS isogenic system based on the generated data.

Our study detected abnormal morphology characterised by shorter and thicker neurites with less branching and terminal points. Importantly, the same phenotype was observed in the human brain. Additionally, T21 spheroids and hCSAs were significantly smaller compared to D21, with fewer proliferating cells and a higher proportion of dying cells throughout differentiation. Finally, these morphological changes were supported by abnormal mitochondrial morphology and synaptogenesis throughout *in vitro* differentiation.

Overall, our study demonstrated that T21 caused abnormal neuronal morphology in both 2D neurons and hCSAs, as well as in the human brain. This is important, as it opens experimental opportunities to mechanistically and genetically model these individual cellular phenotypes *in vitro*, potentially assigning them to specific actions of overdosed HSA21-encoded proteins. Such studies could be a vehicle to arrive at specific therapeutic approaches to ameliorate these aberrations.

## METHODS

### Isogenic iPSCs

In this study, isogenic iPSCs, three disomic (D21: C3, C7 and C9) and three trisomic (T21: C5, C6 and C13) lines were used [3]. Two additional lines were generated by the Nižetić group: one disomic (D21C3GFP) and one trisomic (T21C5GFP) into which a green fluorescent protein was stably introduced using a lentiviral construct [30]. Cultures were tested and maintained mycoplasma free (Cat. No. 20-700-20). In brief, iPSCs were cultured in 6-well plates coated with Geltrex (Cat. No. A14133-02) using Essential 8 Medium (E8) (Supplementary Table 2). For passaging iPSCs colonies, cells were treated with ReLeSR (Cat. No. 100-0484), resuspended in E8 and transferred to new 6-well plates.

### Generation of isogenic NSCs from iPSCs

NSCs were generated from isogenic iPSCs following a previously published GIBCO protocol [65]. Briefly, iPSCs were maintained in E8 medium under standard culture conditions until reaching approximately 20% confluency, after which the medium was replaced with Neural Induction Medium (NIM) (Supplementary Table 3) to induce NSCs differentiation over seven days. At DIV7, NSCs were passaged, maintained and expanded in Neural Expansion Medium (NEM) (Supplementary Table 4). The NSCs were validated by immunofluorescence using specific primary antibodies listed in Supplementary Table 10 and secondary antibodies listed in Supplementary Table 11. Validated NSCs were used for neuronal differentiation. For a detailed protocol, please refer to the Supplementary document.

### Differentiation of isogenic neurons from NSCs

For the neuronal differentiation, isogenic NSCs between passages P5-10 were used. NSCs were first incubated in Accutase (Cat. No. A11105-01) at 37 °C for 4 minutes and dissociated into single cell suspension. Sterile coverslips placed in 24-well plates as well as 6-well plates were coated with Geltrex for 1 hour at 37 °C. Subsequently, 2×10⁵ isogenic NSCs were seeded onto the coverslips, and 1×10⁶ cells were seeded into the 6-well plates in NEM for the next 24 hours. On the following day, half the volume of NEM was replaced with BrainPhys Medium (Supplementary Table 5). BrainPhys Medium was then half media changed twice a week, and neurons were differentiated until DIV60. For IF, WB and qPCR neurons were harvested at following time points: DIV14, DIV40 and DIV60. For a detailed protocol, please refer to the Supplementary document.

### Generation of isogenic hCSs and hStrSs

The isogenic hCSs and hStrSs were generated following a previously published protocol [32] with the following changes. To generate 3D spheroids iPSCs were incubated with Accutase at 37 °C for 4 minutes and dissociated into single cells. E8 Medium at double the volume of Accutase was added to the cells to neutralize the enzyme and a single cell suspension were generated by triturating. The cell suspension was then centrifuged to remove the Accutase after which the pellet was resuspended in E8 Medium supplemented with Penicillin-Streptomycin and 10 µM ROCK inhibitor Y27632 (Cat. No. Y0503). In total, 10,000 cells were seeded per well in 96-well ultra-low-attachment round bottom plate (Cat. No. 7007). The plate was then centrifuged at 200 g for 2 minutes and incubated overnight at 37 °C with 5% CO2. On day 2, E8 Medium was replaced with new Essential 6 Medium (E6) (Supplementary Table 6) which was changed daily until DIV6. To generate hStrSs, at DIV6 the medium was replaced with Neural Differentiation Medium for hStrSs (Supplementary Table 7). To generate hCSs, at DIV6 the medium was replaced with the Neural Differentiation Medium for hCSs (Supplementary Table 8). The culture medium was replaced daily until the point of fusion into assembloids (DIV46). At DIV46 spheroids were transferred into 24-well Clear Flat Bottom Ultra-Low Attachment Well Plates (Cat. No. 3473) in Assembloids Differentiation Medium (Supplementary Table 9). From this stage onward, full media change was performed twice per week. For a detailed protocol, please refer to the Supplementary document.

### Generation of isogenic hCSAs

To generate isogenic hCSAs, hCSs and hStrSs were first derived separately and fused by placing them to the bottom of 1.5 ml tubes and maintained in an incubator for 4 days to facilitate spontaneous fusion. The medium was carefully changed on day 2. By day 4, successfully formed hCSAs were gently transfered into 24-well Clear Flat Bottom Ultra-Low Attachment Well Plates in neural medium as described. Assembloid fusion was performed at DIV46 of differentiation. For a detailed protocol, please refer to the Supplementary document.

### Human brain samples

PFA-fixed, paraffin-embedded human anonymized *post-mortem* brain samples were obtained from the Zagreb Neuroembryological Collection of the Croatian Institute for Brain Research (CIBR) [33]. A detailed list of all samples used in this study is provided in Supplementary Table 1. Slices were cut at 5-10 μm thickness and stained [39] using primary and secondary antibodies listed in Supplementary Tables 10-11.

### Fluorescence *in-situ* hybridisation (FISH)

The number of chromosomes present in isogenic iPSCs was verified by FISH following previously described protocols [39, 66], using the XA 13/18/21 Probe (D-5607-100-TC, MetaSystems Probes). The spots that indicate chromosome 21 were labelled as red signal, whereas the spots that indicate chromosome 13 were labelled as green signal. Over 500 nuclei from eight different Z-stacks were scored for the number of fluorescent signals present and based in this count, nuclei were assigned to four different categories as follows: “one signal”, “two signals”, “three signals” and “> three signals”. Damaged nuclei or nuclei overlapping with neighbouring were not included in scoring.

### Cryopreservation and immunofluorescence (IF)

2D cultures (iPSCs, NSCs and neurons) were fixed in 4% PFA for 15 minutes, rinsed 3 times in PBS and stored at 4 °C while 3D cultures (hStrSs, hCSs, and hCSAs) were fixed in 4% PFA overnight at 4 °C. On the next day, tissues were rinsed 3 times in PBS and transferred to 10% sucrose-PBS at 4 °C overnight followed by 30% sucrose–PBS for 2–3 days until the tissue sank in the solution. Subsequently, they were embedded in Tissue-Tek OCT Compound (Cat. No. 1231400013). For the IF staining, 20 µm-thick sections were cut using a Leica Cryostat (Cat. No. CM 1850). IF was performed as previously reported [28, 39]. Briefly, samples were blocked and permeabilised for 1 hour at room temperature (3% goat serum and 0.2% Triton X-100 in PBS). Samples were subsequently incubated overnight at 4 °C with primary antibodies listed in Supplementary Table 10. The next day, samples were rinsed 3 times in PBS and then incubated with secondary antibodies listed in Supplementary Table 11 for 2 hours in the dark at room temperature. Following washes, nuclei were counterstained with DAPI (D9542).

### Clearing and 3D staining of hCSAs

To optically clear isogenic hCSAs, we used previously published iDISCO+ protocol [67] with the following changes. Briefly, hCSAs at DIV70, DIV100 and DIV130 were fixed with 4% PFA. The next day, hCSAs were rinsed twice with PBS and were dehydrated in an ascending series of MeOH (20%, 40%, 60%, 80% and 100% MeOH for 1 hour each) and left overnight in 66% Dichloromethane (DCM)/MeOH for lipid removal. The next day, hCSAs were rinsed twice with MeOH and left overnight in H_2_O_2_/MeOH solution (5 volume parts of MeOH + one volume part of 30% H_2_O_2_) for depigmentation. The following day, hCSAs were rehydrated in descending series of MeOH (100%, 80%, 60%, 40% and 20% MeOH for 1 hour each), incubated in permeabilization and blocking solution (3% goat serum and 0.2% Triton X-100 in PBS) for 6 hours at room temperature and then stained with primary antibodies listed in Supplementary Table 10 in PBS containing 0.2% Triton X-100 and 3% goat serum at 37 °C for 3 days. hCSAs were subsequently rinsed three times with PBS for 3 hours and then incubated with secondary antibodies (Supplementary Table 11) in PBS containing 0.2% Triton X-100 at 37 °C for 3 days. hCSAs were subsequently rinsed three times with PBS for 2 hours and embedded in 1% Agarose (Cat. No. R0491). Agarose blocks containing hCSAs were dehydrated as described above, cleared in Dibenzyl ether (DBE) and imaged on Ultramicroscope II Super Plan configuration (Miltenyi Biotech). All steps were performed on orbital shaker.

### Transmission electron microscopy (TEM)

hCSAs for ultrastructural analysis by electron microscopy were fixed in McDowell’s fixative, followed by postfixation in 2% osmium tetroxide, contrasting with 3% uranyl acetate, dehydration through a graded acetone series (30-100%), and embedding in synthetic resin Durcupan (Cat. No. 44610). Semithin sections (0.5-1 µm) were sliced using an ultramicrotome, stained with toluidine blue, and examined under a light microscope to identify regions of interest and assess overall tissue morphology. A portion of the tissue was further sectioned into ultrathin sections (50-100 nm) using an ultramicrotome and mounted on copper grids. The grids were placed onto the electron microscope holder (JEOL 1400), and ultrastructural tissue analysis was performed. Tissue regions of interest were imaged using a high-resolution digital camera integrated into the microscope and processed with iTEM software (Olympus Soft Imaging Solutions GmbH, Münster, Germany).

### Scanning electron microscopy (SEM)

Spheroids and hCSAs used in this study were prepared for SEM analyses following previously published protocol [68] with our modifications. Briefly, spheroids and hCSAs were dehydrated in an ascending series of ethanol (70-96%, each 5 min), followed by a solution of 96% ethanol with 100% acetone (1:1) and pure 100% acetone (each 5 min). Following dehydration, samples were dried in a critical point dryer using CO_2_ (Critical Point Dryer K850, LEICA, Wetzlar, Germany). Dried samples were mounted on aluminium stubs and sputtered with a 10 nm gold coat in Sputter Coater (S150B, Edwards, England). Observations and photo documentation of the samples were conducted using an SEM EVO 10 (ZEISS, Germany) at an accelerating voltage of 15 kV.

### Western blotting (WB)

Whole-cell lysates of neurons, spheroids and hCSAs were separated in a NuPAGE™ 4-12% Bis-Tris Gel (Cat. No. NP0323BOX) or NuPAGE™ 10% Bis-Tris Gel (Cat. No. NP0303BOX) and transferred to a nitrocellulose membrane according to the manufacturer’s protocols (Bio-Rad). Following a 60 minutes incubation in 5% non-fat milk (Cat. No. 70166-500G) in TBS-T at room temperature the membrane was incubated with primary and secondary antibodies listed in Supplementary Tables 10-11. The bands were visualized with SuperSignal™ West Femto Maximum Sensitivity Substrate (Cat. No. 34095) while quantification was carried out using BioRad software and density was calculated by Fiji-win 64 software using “gel analysis” option. All signals were normalised to corresponding β-actin loading control for all samples. Uncropped blots were provided in Supplementary Document (Original Western Blots).

### Quantitative polymerase chain reaction (qPCR)

Total RNA from hCSs, hStrSs, hCSAs and neurons was isolated using the RNeasy Mini kit (Cat. No. 74104), cDNA was prepared by reverse transcription using high-capacity RNA-to-cDNA Kit (Cat. No. 4368814). qPCR was performed using the PowerTrack^TM^ SYBR Green Master Mix for qPCR (Cat. No. A46110) on a 7500 Real-Time PCR system. qRT-PCR primer design and in-silico validation for each target gene, the transcript was acquired using NCBI Gene bank (https://www.ncbi.nlm.nih.gov/gene/). Three primer pairs were designed using NCBI Primer-BLAST (https://www.ncbi.nlm.nih.gov/tools/primer-blast) with parameters optimized for quantitative real-time PCR: amplicon size 70-180 bp, primer melting temperature (Tm) 58-62 °C, and GC content 40-60%. Primers were screened for specificity against the relevant genome and transcriptome, and the amplicon size was confirmed using the UCSC Genome Browser In-Silico PCR tool. Primers were synthesized commercially by SIGMA ALDRICH. Primers used in this study are listed in (Supplementary Table 12). Gene expression was calculated using the ΔCT method.

### Flow cytometry

For flow cytometry analysis isogenic iPSCs, isogenic spheroids at DIV1 and DIV5, and hCSAs at DIV70 were used. iPSCs were rinsed 2 times in Dulbecco’s PBS (D-PBS) and incubated with Accutase at 37 °C for 4 minutes. D-PBS at double the volume of Accutase was added to the cells and a single cell suspension was generated by triturating. To remove any remaining cell aggregates the suspension was passed through a 40 µm cell strainer. Cells were centrifuged to remove the Accutase and cell pellet were resuspended in D-PBS. Spheroids and hCSAs were dissociated using a combination of enzymatic digestion and mechanical dissociation. In total, a pool of five to six spheroids or hCSAs were used for both, D21 and T21 samples which were transferred into 15 mL tubes. The samples were rinsed two times in D-PBS to completely remove remaining culture media and 1 mL of prewarmed Accutase was added. The hCSAs were incubated at 37 °C for 5 minutes and then mechanically dissociated by gentle pipette trituration. The incubation and the pipette trituration step was repeated once more to ensure complete dissociation into single cells. The samples were filtered through 40 µm cell strainer, centrifuged at 300 g for 4 minutes and resuspend in D-PBS.

Single cell suspensions were used for flow cytometry analysis on an instrument BD FACSAria IIu (BD Biosciences, Franklin Lakes, NJ, USA). For the analysis of cells derived from spheroids, the cell viability was determined using DAPI. For the analysis of cells derived from hCSAs, the percentage of GFP positive cells was determined by setting gates according to GFP negative control samples. In addition, to visualize the active mitochondria, cells were incubated with MitoTracker probes (Cat. No. M7512) according to the manufacturer’s instructions. Briefly, NSCs were cultured, incubated with MitoTracker dyes (final concentration 1 µM) for 1 hour and prepared for flow cytometry as described above. Gates were set to determine the proportion of cells with strong MitoTracker fluorescence, according to unlabelled samples. Data were analysed in FlowJo software (BD Biosciences).

### Image analysis

Confocal images were captured by Olympus FV 3000 microscope and further analysed using Imaris 9.9.1 - Neuroscience module software (BITPLANE, An Oxford Instruments Co., Zurich, Switzerland). Quantification was performed blinded to the genotype, on 5-8 full Z-stacks images/staining/time point, using “surface”, “spot” and “filament” options as well as Pearson’s coefficient of colocalization. For quantification of specific markers total fluorescence intensity of positive signals was normalised to the total fluorescence intensity of DAPI as a nuclear dye or total fluorescence intensity of MAP2 as a pan-neuronal marker. The percentage of nuclear markers positive cells were also counted using the “spot” option and normalised with number of DAPI positive nuclei. For the Filament analysis, the body of the neurons was defined as the point of origin, following which the software autonomously computed subsequent options without human intervention: “Filament Diameter”, “Filament Length”, “Filament branches”, “Filament No. branching points”, “Filament No. Terminal Points” and “Sholl Intersections” [69]. The “Filament diameter” was defined as the mean value of the diameters of all processes, “Filament length” was defined as a length of all processes per field of view, “Filament branches” was defined as the total number of branches per field of view, “Filament No. branching points” was defined as a number of branching points per field of view, “Filament No. Terminal Points” was defined as a number of terminal points per field of view and “No. of Sholl Intersections” was defined as the number of process intersections (branches) on concentric spheres (1.0 µm step). Pearson’s coefficient of colocalization is defined as corelation between two channels and were calculated automatically. Brightfield images were captured by EVOS FL Auto Imaging System and further analysed using Lusca software [70].

### Statistics

Data were analysed using GraphPad Prism (v. 8.4.2.). All data are presented as mean ± standard error of the mean (SEM), unless otherwise indicated. Distribution of the raw data was tested for normality of distribution. Statistical analyses were performed using a two-tailed Student’s T test when comparing single pair of samples and one- or two-way analysis of variance (ANOVA) followed by Bonferroni’s multiple comparisons test for experiments involved multiple groups or variable. Statistical significance was defined at the levels of p<0.05 (*), p<0.01 (**), p<0.001 (***), and p<0.0001 (****). Blinding was performed for imaging experiments and analyses. P values were shown in Supplementary Table 13.

## Data availability

All data generated or analysed during this study are available within the article and Supplementary Files. Additional supporting data are available from the corresponding authors upon reasonable request.

## Acknowledgements

Human brain tissue collection procedures complied with the ethical principles outlined in the 2000 Declaration of Helsinki. Informed consent was obtained from all patients. The samples were obtained from the Zagreb Neuroembryological Collection of the Croatian Institute for Brain Research (CIBR) [33], with ethical approval: (1) Internal Review Bord of the Ethical Committee of the Faculty of Veterinary Medicine (640-01/22-02/10; 251/61-01/139-22-50) and (2) Internal Review Bord of the Ethical Committee of the School of Medicine University of Zagreb (651-01/23-02/01; 251-59-10106-23-111/226).

We gratefully acknowledge Prof. Mario Vukšić from the School of Medicine Imaging Facility for his valuable assistance. The authors also thank Danica Budinščak, Leonarda Grandverger and Sandra Grgić for its technical assistance. We thank Ms. Tara Adelaide Rutherford for proofreading the manuscript.

## Author contributions

AP and IA contributed to the concept and design, the acquisition, analysis and interpretation, and drafting of manuscript. GG, AM, DN – isogenic iPSCs, NSCs, Neuronal and Cerebral Organoid technology, primer design, analysis and data interpretation; DG – Flow cytometry; MH – TEM analysis; KSS, HJ – SEM analysis; IŠ – Lusca software and Light-sheet microscope; ŽK – human brain tissue; DM – iPSCs technology, laboratory equipment, consumables and software analysis. All authors contributed to the writing of the manuscript.

## References

1. Antonarakis SE, Skotko BG, Rafii MS, Strydom A, Pape SE, Bianchi DW, et al. Down syndrome. Nat Rev Dis Primers. 2020; 6:9. https://pmc.ncbi.nlm.nih.gov/articles/PMC8428796/.

2. Wiseman FK, Al-Janabi T, Hardy J, Karmiloff-Smith A, Nizetic D, Tybulewicz VLJ, et al. A genetic cause of Alzheimer disease: mechanistic insights from Down syndrome. Nat Rev Neurosci. 2015; 16:564–74. https://pmc.ncbi.nlm.nih.gov/articles/PMC4678594/.

3. Murray A Letourneau A, Canzonetta C, Stathaki E, Gimelli S, Sloan-Bena F, et al. Brief report: isogenic induced pluripotent stem cell lines from an adult with mosaic down syndrome model accelerated neuronal ageing and neurodegeneration. Stem Cells. 2015; 33:2077–84. https://pmc.ncbi.nlm.nih.gov/articles/PMC4737213/.

4. Papavassiliou P, York TP, Gursoy N, Hill G, Vanner Nicely L, Sundaram U, et al. The Phenotype of Persons Having Mosaicism for Trisomy 21/Down Syndrome Reflects the Percentage of Trisomic Cells Present in Different Tissues. Am J Med Genet A. 2009; 0:573–583. https://onlinelibrary.wiley.com/doi/10.1002/ajmg.a.32729.

5. Papavassiliou P, Charalsawadi C, Rafferty K, Jackson-Cook C. Mosaicism for trisomy 21: A review. Am J Med Genet A. 2015; 167A:26–39. https://onlinelibrary.wiley.com/doi/10.1002/ajmg.a.36861.

6. Gafson AR, Barthélemy NR, Bomont P, Carare RO, Durham HD, Julien JP, et al. Neurofilaments: neurobiological foundations for biomarker applications. Brain. 2020; 143:1975–1998. https://pmc.ncbi.nlm.nih.gov/articles/PMC7363489/.

7. Yuan A, Sershen H, Veeranna, Basavarajappa BS, Kumar A, Hashim A, et al. Functions of neurofilaments in synapses. Mol Psychiatry. 2015; 20:915. https://www.nature.com/articles/mp201599.

8. Yuan A, Nixon RA. Posttranscriptional regulation of neurofilament proteins and tau in health and disease. Brain Res Bull. 2023; 192:115–127. https://www.sciencedirect.com/science/article/pii/S0361923022002957?via%3Dihub.

9. Ratnam J, Teichberg VI. Neurofilament-light increases the cell surface expression of the N-methyl-D-aspartate receptor and prevents its ubiquitination. J Neurochem. 2005; 92:878–85. https://onlinelibrary.wiley.com/doi/10.1111/j.1471-4159.2004.02936.x.

10. Yuan A, Veeranna, Sershen H, Basavarajappa BS, Smiley JF, Hashim A, et al. Neurofilament light interaction with GluN1 modulates neurotransmission and schizophrenia-associated behaviors. Transl Psychiatry. 2018; 8:167. https://www.nature.com/articles/s41398-018-0194-7.

11. Schulz JM, Knoflach F, Hernandez MC, Bischofberger J. Enhanced Dendritic Inhibition and Impaired NMDAR Activation in a Mouse Model of Down Syndrome. J Neurosci. 2019; 39:5210–5221. https://www.jneurosci.org/content/39/26/5210.long.

12. Bocquet A, Berges R, Frank R, Robert P, Peterson AC, Eyer J. Neurofilaments bind tubulin and modulate its polymerization. J Neurosci. 2009; 29:11043–54. https://www.jneurosci.org/content/29/35/11043.long.

13. Yau KW, Schätzle P, Tortosa E, Pagès S, Holtmaat A, Kapitein LC, et al. Dendrites In Vitro and In Vivo Contain Microtubules of Opposite Polarity and Axon Formation Correlates with Uniform Plus-End-Out Microtubule Orientation. J Neurosci. 2016; 36:1071–85. https://www.jneurosci.org/lookup/pmidlookup?view=long&pmid=26818498.

14. Currey L, Harvey T, Pelenyi A, Piper M, Thor S. Mechanisms of brain overgrowth in autism spectrum disorder with macrocephaly. Front Neurosci. 2025; 19:1586550. https://www.frontiersin.org/journals/neuroscience/articles/10.3389/fnins.2025.1586550/full.

15. Schwartz ML, Shneidman PS, Bruce J, Schlaepfer WW. Stabilization of neurofilament transcripts during postnatal development. Brain Res Mol Brain Res. 1994; 27:215–20. https://www.sciencedirect.com/science/article/abs/pii/0169328X94900035?via%3Dihub.

16. Schwartz ML, Bruce J, Shneidman PS, Schlaepfer WW. Deletion of 3’-untranslated region alters the level of mRNA expression of a neurofilament light subunit transgene. J Biol Chem. 1995; 270:26364–9. https://www.sciencedirect.com/science/article/pii/S0021925818924877?via%3Dihub.

17. Hannan SB, Alić I, Murray A, Kwon J, Mortensen M, Kang HJ, et al. Synaptic and intrinsic membrane defects disrupt early neural network dynamics. Nat. Commun. 2026; 17: 1287. https://www.nature.com/articles/s41467-025-68048-x

18. Rangaraju V, Lewis TL, Hirabayashi Y, Bergami M, Motori E, Cartoni R, et al. Pleiotropic Mitochondria: The Influence of Mitochondria on Neuronal Development and Disease. J Neurosci. 2019; 39:8200–8208. https://www.jneurosci.org/content/39/42/8200.long.

19. Keller N, Christensen TA, Wanberg EJ, Salisbury JL, Trushina E. Neuroprotective mitochondria targeted small molecule restores synapses and the distribution of synaptic mitochondria in the hippocampus of APP/PS1 mice. Sci Rep. 2025; 15:6528. https://www.nature.com/articles/s41598-025-90925-0.

20. Tao K, Matsuki N, Koyama R. AMP-activated protein kinase mediates activity-dependent axon branching by recruiting mitochondria to axon. Dev Neurobiol. 2014; 74:557–73. https://onlinelibrary.wiley.com/doi/10.1002/dneu.22149.

21. Courchet J, Lewis TL, Lee S, Courchet V, Liou DY, Aizawa S, et al. Terminal axon branching is regulated by the LKB1-NUAK1 kinase pathway via presynaptic mitochondrial capture. Cell. 2013; 153:1510–25. https://www.sciencedirect.com/science/article/pii/S0092867413005886?via%3Dihub.

22. Spillane M, Ketschek A, Merianda TT, Twiss JL, Gallo G. Mitochondria coordinate sites of axon branching through localized intra-axonal protein synthesis. Cell Rep. 2013; 5:1564–75. https://www.sciencedirect.com/science/article/pii/S221112471300689X?via%3Dihub.

23. Bahn S, Mimmack M, Ryan M, Caldwell MA, Jauniaux E, Starkey M, et al. Neuronal target genes of the neuron-restrictive silencer factor in neurospheres derived from fetuses with Down’s syndrome: a gene expression study. Lancet. 2002; 359:310–5. https://www.sciencedirect.com/science/article/pii/S0140673602074974?via%3Dihub.

24. Khalil M, Teunissen CH, Otto M, Piehl F, Sormani MP, Gattringer T, et al. Neurofilaments as biomarkers in neurological disorders. Nat Rev Neurol. 2018; 14:577–589. https://www.nature.com/articles/s41582-018-0058-z.

25. Weick JP, Held DL, Bonadurer GF 3rd, Doers ME, Liu Y, Maguire C, et al. Deficits in human trisomy 21 iPSCs and neurons. Proc Natl Acad Sci U S A. 2013; 110:9962-7. https://pmc.ncbi.nlm.nih.gov/articles/PMC3683748/.

26. Giffin-Rao Y, Sheng J, Strand B, Xu K, Huang L, Medo M, et al. Altered patterning of trisomy 21 interneuron progenitors. Stem Cell Reports. 2022; 17:1366–1379. https://www.sciencedirect.com/science/article/pii/S2213671122002090?via%3Dihub.

27. Huo HQ, Qu ZY, Yuan F, Ma L, Yao L, Xu M, et al. Modeling Down Syndrome with Patient iPSCs Reveals Cellular and Migration Deficits of GABAergic Neurons. Stem Cell Reports. 2018; 10:1251–1266. https://www.sciencedirect.com/science/article/pii/S2213671118300699?via%3Dihub.

28. Murray A, Gough G, Cindriæ A, Vuèkoviæ F, Koschut D, Borelli V, et al. Dose imbalance of DYRK1A kinase causes systemic progeroid status in Down syndrome by increasing the un-repaired DNA damage and reducing LaminB1 levels. EBioMedicine. 2023; 94:104692. https://www.sciencedirect.com/science/article/pii/S2352396423002578?via%3Dihub.

29. Canzonetta C, Mulligan C, Deutsch S, Ruf S, O’Doherty A, Lyle R, et al. DYRK1A-dosage imbalance perturbs NRSF/REST levels, deregulating pluripotency and embryonic stem cell fate in Down syndrome. Am J Hum Genet. 2008; 83:388–400. https://www.sciencedirect.com/science/article/pii/S0002929708004503?via%3Dihub.

30. Gough, G. The genetic dissection of trisomy 21 and partial trisomy 21 cellular pathologies using induced pluripotent stem cells. 2021. doi:10.32657/10356/153265.

31. Miura Y, Li MY, Birey F, Ikeda K, Revah O, Thete MV, et al. Generation of human striatal organoids and cortico-striatal assembloids from human pluripotent stem cells. Nat Biotechnol. 2020; 38:1421–1430. https://www.nature.com/articles/s41587-020-00763-w.

32. Miura Y, Li MY, Revah O, Yoon SJ, Narazaki G, Pașca SP. Engineering brain assembloids to interrogate human neural circuits. Nat Protoc. 2022; 17:15–35. https://www.nature.com/articles/s41596-021-00632-z.

33. Hrabač P, Bosak A, Vukšić M, Judaš M, Kostović I, Krsnik Ž. The Zagreb Collection of human brains: entering the virtual world. Croat Med J. 2018; 59:283–287. https://www.cmj.hr/2018/59/6/30610769.htm.

34. Busciglio J, Yankner BA. Apoptosis and increased generation of reactive oxygen species in Down’s syndrome neurons in vitro. Nature. 1995; 378:776–9. https://www.nature.com/articles/378776a0.

35. Bhattacharyya A, McMillan E, Chen SI, Wallace K, Svendsen CN. A critical period in cortical interneuron neurogenesis in down syndrome revealed by human neural progenitor cells. Dev Neurosci. 2009; 31:497–510. https://karger.com/dne/article/31/6/497/107302/A-Critical-Period-in-Cortical-Interneuron.

36. Olmos-Serrano JL, Kang HJ, Tyler WA, Silbereis JC, Cheng F, Zhu Y, et al. Down Syndrome Developmental Brain Transcriptome Reveals Defective Oligodendrocyte Differentiation and Myelination. Neuron. 2016; 89:1208–1222. https://pmc.ncbi.nlm.nih.gov/articles/PMC4795969/.

37. Agrawal M, Kirkise N, Rygel K, Sahoo PK, Vincent CJ, Joyner D, et al. Restoring DSCAM expression rescues neuronal morphology and axon guidance deficits in Down syndrome. bioRxiv [Preprint]. 2025. https://www.biorxiv.org/content/10.1101/2025.07.25.666804v1.

38. Blackburn E, Birsa N, Lopes AT, Fisher EMC, Lazo OM, Schiavo G. Impaired BDNF-TrkB trafficking and signalling in Down syndrome basal forebrain neurons. Cell Death Dis. 2026; 17:214. https://www.nature.com/articles/s41419-026-08464-z

39. Alić I, Goh PA, Murray A, Portelius E, Gkanatsiou E, Gough G, et al. Patient-specific Alzheimer-like pathology in trisomy 21 cerebral organoids reveals BACE2 as a gene dose-sensitive AD suppressor in human brain. Mol Psychiatry. 2021; 26: 5766–5788. https://www.nature.com/articles/s41380-020-0806-5.

40. Guidi S, Bonasoni P, Ceccarelli C, Santini D, Gualtieri F, Ciani E, et al. Neurogenesis impairment and increased cell death reduce total neuron number in the hippocampal region of fetuses with Down syndrome. Brain Pathol. 2008; 18:180–97. https://onlinelibrary.wiley.com/doi/10.1111/j.1750-3639.2007.00113.x.

41. Fernández-Blanco A, González-Arias C, Sierra C, Zamora-Moratalla A, Perea G, Dierssen M. Astrocytopathy Is Associated with CA1 Synaptic Dysfunction in a Mouse Model of Down Syndrome. Cells. 2025; 14:1332. https://www.mdpi.com/2073-4409/14/17/1332.

42. Herrera F, Chen Q, Fischer WH, Maher P, Schubert DR. Synaptojanin-1 plays a key role in astrogliogenesis: possible relevance for Down’s syndrome. Cell Death Differ. 2009; 16:910–20. https://www.nature.com/articles/cdd200924.

43. Verkhratsky A, Butt A, Li B, Illes P, Zorec R, Semyanov A, et al. Astrocytes in human central nervous system diseases: a frontier for new therapies. Signal Transduct Target Ther. 2023; 8:396. https://pmc.ncbi.nlm.nih.gov/articles/PMC10570367/.

44. van Asperen JV, Kotaich F, Caillol D, Bomont P. Neurofilaments: Novel findings and future challenges. Curr Opin Cell Biol. 2024; 87:102326. https://www.sciencedirect.com/science/article/pii/S095506742400005X?via%3Dihub.

45. Sheng ZH, Cai Q. Mitochondrial transport in neurons: Impact on synaptic homeostasis and neurodegeneration. Nat Rev Neurosci. 2012; 13:77–93. https://pmc.ncbi.nlm.nih.gov/articles/PMC4962561/.

46. Prutton KM, Marentette JO, Maclean KN, Roede JR. Characterization of mitochondrial and metabolic alterations induced by trisomy 21 during neural differentiation. Free Radic Biol Med. 2023; 196:11–21. https://pmc.ncbi.nlm.nih.gov/articles/PMC9898228/.

47. Xu L, Huo HQ, Lu KQ, Tang XY, Hong Y, Han X, et al. Abnormal mitochondria in Down syndrome iPSC-derived GABAergic interneurons and organoids. Biochim Biophys Acta Mol Basis Dis. 2022; 1868:166388. https://www.sciencedirect.com/science/article/pii/S0925443922000515?via%3Dihub.

48. Chaklader M, Souza Bomfim GH, Jeju N, Duan Y, Niemeyer BF, Espinosa JM, et al. Elevated mitochondrial metabolism in Down syndrome iPSCs reduces commitment to neuroectoderm. bioRxiv 2025. https://www.biorxiv.org/content/10.1101/2025.05.09.653211v1.

49. Lana-Elola E, Aoidi R, Llorian M, Gibbins D, Buechsenschuetz C, Bussi C, et al. Increased dosage of DYRK1A leads to congenital heart defects in a mouse model of Down syndrome. Sci Transl Med. 2024; 16:eadd6883. https://pmc.ncbi.nlm.nih.gov/articles/PMC7615651/.

50. Izzo A, Mollo N, Cicatiello R, Genesio R, Paladino S, Conti A, et al. Mitochondrial Abnormalities in Down Syndrome: Pathogenesis, Effects and Therapeutic Approaches. Adv. Res. Down Syndr. 2018; 109–38. https://www.intechopen.com/chapters/57817

51. Duarte FV, Ciampi D, Duarte CB. Mitochondria as central hubs in synaptic modulation. Cell Mol Life Sci. 2023; 80:173. https://link.springer.com/article/10.1007/s00018-023-04814-8.

52. DiFilippo A, Jonaitis E, Makuch R, Gambetti B, Fleming V, Ennis G, et al. Measurement of synaptic density in Down syndrome using PET imaging: a pilot study. Sci Rep. 2024; 14:4676. https://www.nature.com/articles/s41598-024-54669-7.

53. Real R, Peter M, Trabalza A, Khan S, Smith MA, Dopp J, et al. In vivo modeling of human neuron dynamics and Down syndrome. Science. 2018; 362:eaau1810. https://www.science.org/doi/10.1126/science.aau1810.

54. Peter M, Real R, Strano A, Robinson HPC, Smith MA, Barnes SJ, et al. Trisomy 21 impairs synchronized activity and connectivity in developing human down syndrome cortical excitatory neuron networks. bioRxiv 2024. https://www.biorxiv.org/content/10.1101/2024.12.27.630530v2.

55. Hannan SB, Lana-Elola E, Watson-Scales S, Fisher EMC, Tybulewicz VLJ, Smart TG. Hippocampal circuit-specific enhancement of GABA-inhibition caused by discrete gene regions in a Down syndrome model. bioRxiv 2025. https://www.biorxiv.org/content/10.1101/2025.07.21.665861v1.

56. Botte A, Laine J, Xicota L, Heiligenstein X, Fontaine G, Kasri A, et al. Ultrastructural and dynamic studies of the endosomal compartment in Down syndrome. Acta Neuropathol. Commun. 2020; 8:89. https://link.springer.com/article/10.1186/ s40478-020-00956-z

57. Tang XY, Xu L, Wang J, Hong Y, Wang Y, Zhu Q, et al. DSCAM/PAK1 pathway suppression reverses neurogenesis deficits in iPSC-derived cerebral organoids from patients with down syndrome. J Clin Invest. 2021; 131:e135763. https://www.jci.org/articles/view/135763.

58. Liu H, Huang S, Wang W, Wang H, Huang W, Zhai Z, et al. Migration deficits of the neural crest caused by CXADR triplication in a human Down syndrome stem cell model. Cell Death Dis. 2022; 13:1018. https://www.nature.com/articles/s41419-022-05481-6

59. Pinter JD, Eliez S, Schmitt JE, Capone GT, Reiss AL. Neuroanatomy of Down’s Syndrome: A High-Resolution MRI Study. Am J Psychiatry. 2001; 158:1659–65. https://psychiatryonline.org/doi/10.1176/appi.ajp.158.10.1659?url_ver=Z39.88-2003&rfr_id=ori:rid:crossref.org&rfr_dat=cr_pub%20%200pubmed.

60. Hamner T, Udhnani MD, Osipowicz KZ, Lee NR. Pediatric brain development in Down syndrome: A field in its infancy. J Int Neuropsychol Soc. 2018; 24:966–976. https://pmc.ncbi.nlm.nih.gov/articles/PMC6207466/.

61. Rodrigues M, Nunes J, Figueiredo S, Martins de Campos A, Geraldo AF. Neuroimaging assessment in Down syndrome: a pictorial review. Insights Imaging. 2019; 10:52. https://link.springer.com/article/10.1186/s13244-019-0729-3.

62. Levman J, McCann B, Baumer N, Lam MY, Shiohama T, Cogger L, et al. Structural Magnetic Resonance Imaging-Based Surface Morphometry Analysis of Pediatric Down Syndrome. Biology (Basel). 2024; 13:575. https://www.mdpi.com/2079-7737/13/8/575.

63. Ferrer I, Gullotta F. Down’s syndrome and Alzheimer’s disease: dendritic spine counts in the hippocampus. Acta Neuropathol. 1990; 79:680–5. https://pubmed.ncbi.nlm.nih.gov/2141748/.

64. Shabani K, Pigeon J, Zariouh MBT, Liu T, Saffarian A, Komatsu J, et al. The temporal balance between self-renewal and differentiation of human neural stem cells requires the amyloid precursor protein. Sci Adv. 2023; 9: eadd5002. https://pubmed.ncbi.nlm.nih.gov/37327344/

65. Technologies, L. Induction of Neural Stem Cells from Human Pluripotent Stem Cells Using PSC Neural Induction Medium (MAN0008031 Rev A.0).

66. Solovei I, Cremer M. 3D-FISH on cultured cells combined with immunostaining. Methods Mol Biol. 2010; 659:117–26. https://link.springer.com/protocol/10.1007/978-1-60761-789-1_8.

67. Renier N, Wu Z, Simon DJ, Yang J, Ariel P, Tessier-Lavignei M. iDISCO: a simple, rapid method to immunolabel large tissue samples for volume imaging. Cell. 2014; 159:896–910. https://www.sciencedirect.com/science/article/pii/S0092867414012975?via%3Dihub.

68. Plewa B, Jackowiak H. The three-dimensional analysis of gustatory papillae and its taste buds on the tongue of the wild-living hare (Lepus europaeus), European rabbit (Oryctolagus cuniculus), and domestic rabbit (Oryctolagus cuniculus f. domestica). Ann Anat. 2025: 260:152667. https://www.sciencedirect.com/science/article/pii/S0940960225002948?via%3Dihub.

69. Plećaš A, Kapuralin K, Grandverger L, Mitrečić D, Alić I. Thy1-YFP: an effective tool for single cell tracing from neuronal progenitors to mature functionally active neurons. Cell Death Discov. 2025; 11:18. https://www.nature.com/articles/s41420-025-02297-z.

70. Šimuniæ I, Jagečić D, Isaković J, Dobrivojević Radmilović M, Mitrečić D. Author Correction: Lusca: FIJI (ImageJ) based tool for automated morphological analysis of cellular and subcellular structures. Sci Rep. 2024; 14:9026. https://www.nature.com/articles/s41598-024-59795-w.

