## Supplementary document for "Abnormal neuronal and synaptic morphology in Down syndrome brains reproduces in human isogenic cellular models"

**Supplementary Table 1.** ***Postmortem*** **human tissues.**

| Sample | Genotype | Gestation week/Age |
|---|---|---|
| 1 | Euploid | 23 gw |
| 2 | Euploid | 22/23 gw |
| 3 | Euploid | 18 gw |
| 4 | DS | 20 gw |
| 5 | DS | 21 gw |
| 6 | DS | 18 gw |
| 7 | Euploid | Newborn |
| 8 | Euploid | Newborn |
| 9 | Euploid | Newborn |
| 10 | Euploid | Newborn |
| 11 | DS | Newborn |
| 12 | DS | Newborn |
| 13 | Euploid | 21 years |
| 14 | Euploid | 27 years |
| 15 | DS | 28 years |

**Supplementary Table 2. Essential 8 Medium (E8).**

| Composition | Volume (50 mL total) | Final concentration | Source |
|---|---|---|---|
| Essential 8™ Medium | 48.5 mL | | ThermoFisher SCIENTIFIC, A1517001 |
| Essential 8™ Supplement (50X) | 1 mL | 2 % | ThermoFisher SCIENTIFIC, A1517101 |
| Penicillin–Streptomycin | 500 µL | 1 % | ThermoFisher SCIENTIFIC, 15140-122 |

**Supplementary Table 3. Neural Induction Medium (NIM).**

| Composition | Volume (50 mL total) | Final concentration | Source | Note |
|---|---|---|---|---|
| Neurobasal™ Medium | 48.5 mL | | ThermoFisher SCIENTIFIC, 21103049 | Part of the kit |
| Neural Induction Supplement | 1 mL | 2 % | ThermoFisher SCIENTIFIC, A1647801 | Part of the kit |
| Penicillin–Streptomycin | 500 µL | 1 % | ThermoFisher SCIENTIFIC, 15140-122 | |

**Supplementary Table 4. Neural Expansion Medium (NEM).**

| Composition | Volume (50 mL total) | Final concentration | Source | Note |
|---|---|---|---|---|
| Neurobasal™ Medium | 24.25 mL | | ThermoFisher SCIENTIFIC, 21103049 | Part of the kit |
| Advanced DMEM/F-12 | 24.25 mL | | ThermoFisher SCIENTIFIC, 12634010 | |
| Neural Induction Supplement | 1 mL | 2 % | ThermoFisher SCIENTIFIC, A1647801 | Part of the kit |
| Penicillin–Streptomycin | 500 µL | 1 % | ThermoFisher SCIENTIFIC, 15140-122 | |

**Supplementary Table 5. BrainPhys Medium.**

| Composition | Volume (50 mL total) | Final concentration | Source | Note |
|---|---|---|---|---|
| BrainPhys™ Neuronal Medium Kit | 48 mL | | STEMCELL TECHNOLOGIES, 05793 | Part of the kit |
| N2-A | 500 µL | 1 % | | Part of the kit |
| SM1 | 1 mL | 2 % | | Part of the kit |
| Dibutyryl CAMP | 500 µL | 1mM (1:100) | Sigma-Aldrich, D0627 | |
| BDNF Recombinant Protein | 50 µL | 20 ng/mL (1:1000) | PEPROTECH, 450-02 | |
| GDNF Recombinant Protein | 50 µL | 20 ng/mL (1:1000) | PEPROTECH, 450-10 | |
| Ascorbic Acid | 50 µL | 200 µM | Sigma-Aldrich, 49752 | |
| Penicillin–Streptomycin | 500 µL | 1 % | ThermoFisher SCIENTIFIC, 15140-122 | |

**Supplementary Table 6. Essential 6 Medium (E6).**

| Composition | Volume (50 mL total) | Final concentration | Source | Note |
|---|---|---|---|---|
| Essential 6™ Medium | 49 mL | | ThermoFisher SCIENTIFIC, A1516401 | |
| MEM Non-essential Amino Acid Solution (100×) | 500 µL | 1 % | Sigma-Aldrich, M7145 | |
| Penicillin–Streptomycin | 500 µL | 1 % | ThermoFisher SCIENTIFIC, 15140-122 | |
| Dorsomorphin | 25 µL | 2.5 µM (1:2000) | Sigma-Aldrich, P5499 | Add just before use |
| SB-431542 | 50 µL | 10 µM (1:1000) | TOCRIS, 1614 | Add just before use |

**Supplementary Table 7. Neural Differentiation Medium (NM) for hStrSs.**

| Composition | Volume (50 mL total) | Final concentration | Source | Note |
|---|---|---|---|---|
| Neurobasal™-A Medium | 47.5 mL | | ThermoFisher SCIENTIFIC, 10888022 | |
| B-27™ Supplement (50X), minus vitamin A | 1 mL | 2 % | ThermoFisher SCIENTIFIC, 12587010 | Needed from day 6 to day 46 |
| Penicillin–Streptomycin | 500 µL | 1 % | ThermoFisher SCIENTIFIC, 15140-122 | |
| GlutaMAX™ Supplement | 500 µL | 1 % | ThermoFisher SCIENTIFIC, 35050061 | |
| MEM Non-essential Amino Acid Solution (100×) | 500 µL | 1 % | Sigma-Aldrich, M7145 | |
| IWP-2 | 50 µL | 2.5 µM (1:1000) | Sigma-Aldrich, I0536 | Needed from day 6 to day 22. Add just before use |
| Activin A Recombinant Protein | 50 µL | 50 ng/mL (1:1000) | PEPROTECH, 120-14P | Needed from day 6 to day 22. Add just before use |
| SR11237 | 50 µL | 100 nM (1:1000) | TOCRIS, 3411 | Needed from day 6 to day 22. Add just before use |
| BDNF Recombinant Protein | 50 µL | 20 ng/mL (1:1000) | PEPROTECH, 450-02 | Needed from day 6 to day 22. Add just before use |
| NT-3 Recombinant Protein | 50 µL | 20 ng/ml (1:1000) | PEPROTECH, 450-03 | Needed from day 6 to day 22. Add just before use |
| Ascorbic Acid | 50 µL | 200 µM (1:1000) | Sigma-Aldrich, 49752 | Needed from day 6 to day 22. Add just before use |
| Cis-DHA | 50 µL | 10 µM (1:1000) | Sigma-Aldrich, D2534 | Needed from day 6 to day 22. Add just before use |
| Dibutyryl CAMP | 25 µL | 50 µM (1:2000) | Sigma-Aldrich, D0627 | Needed from day 6 to day 22. Add just before use |
| DAPT | 25 µL | 2.5 µM (1:2000) | STEMCELL TECHNOLOGIES, 72082 | Needed from day 6 to day 22. Add just before use |

**Supplementary Table 8. Neural Differentiation Medium (NM) for hCSs.**

| Composition | Volume (50 mL total) | Final concentration | Source | Note |
|---|---|---|---|---|
| Neurobasal™-A Medium | 47.5 mL | | ThermoFisher SCIENTIFIC, 10888022 | |
| B-27™ Supplement (50X), minus vitamin A | 1 mL | 2 % | ThermoFisher SCIENTIFIC, 12587010 | Needed from day 6 to day 46 |
| Penicillin–Streptomycin | 500 µL | 1 % | ThermoFisher SCIENTIFIC, 15140-122 | |
| GlutaMAX™ Supplement | 500 µL | 1 % | ThermoFisher SCIENTIFIC, 35050061 | |
| MEM Non-essential Amino Acid Solution (100×) | 500 µL | 1 % | Sigma-Aldrich, M7145 | |
| FGF-2 Recombinant Protein | 50 µL | 20 ng/mL (1:1000) | ThermoFisher SCIENTIFIC, PHG0021 | Needed from day 6 to day 22. Add just before use |
| EGF Recombinant Protein | 50 µL | 20 ng/mL (1:1000) | ThermoFisher SCIENTIFIC, PHG0311 | Needed from day 6 to day 22. Add just before use |
| BDNF Recombinant Protein | 50 µL | 20 ng/mL (1:1000) | PEPROTECH, 450-02 | Needed from day 6 to day 22. Add just before use |
| NT-3 Recombinant Protein | 50 µL | 20 ng/mL (1:1000) | PEPROTECH, 450-03 | Needed from day 6 to day 22. Add just before use |
| Ascorbic Acid | 50 µL | 200 µM (1:1000) | Sigma-Aldrich, 49752 | Needed from day 6 to day 22. Add just before use |
| Cis-DHA | 50 µL | 10 µM (1:1000) | Sigma-Aldrich, D2534 | Needed from day 6 to day 22. Add just before use |
| Dibutyryl CAMP | 25 µL | 50 µM (1:2000) | Sigma-Aldrich, D0627 | Needed from day 6 to day 22. Add just before use |

**Supplementary Table 9. Assembloids Differentiation Medium.**

| Composition | Volume (50 mL total) | Final concentration | Source | Note |
|---|---|---|---|---|
| Neurobasal™-A Medium | 48 mL | | ThermoFisher SCIENTIFIC, 10888022 | |
| B-27™ Supplement (50X) | 1 mL | 2 % | ThermoFisher SCIENTIFIC, 17504044 | |
| Penicillin–Streptomycin | 500 µL | 1 % | ThermoFisher SCIENTIFIC, 15140-122 | |
| GlutaMAX™ Supplement | 500 µL | 1 % | ThermoFisher SCIENTIFIC, 35050061 | |

**Supplementary Table 10. Primary antibodies used in this study.**

| Primary antibody | Clone | Isotype | Source | Dilution | | |
|---|---|---|---|---|---|---|
| | | | | Cells | Tissues | WB |
| AIF | D39D2 | Rabbit IgG monoclonal | Cell Signaling (5318S) | 1:400 | | 1:5000 |
| β-actin | AC-15 | Mouse monoclonal | Sigma-Aldrich (A5441) | | | 1:60000 |
| Cleaved Caspase-3 | ASP 175 | Rabbit IgG polyclonal | Cell Signaling (9661S) | | 1:400 | |
| Ctip2 | 25B6 | Rat IgG2a monoclonal | Abcam (ab18465) | 1:200 | 1:200 | |
| Doublecortin (DCX) | | Goat IgG polyclonal | Abcam (ab223435) | 1:1000 | | |
| Gephyrin | EPR12650 | Rabbit IgG monoclonal | Abcam (ab181382) | 1:200 | 1:200 | |
| GFAP | 2.2B10 | Rat IgG2a monoclonal | ThermoFisher SCIENTIFIC (13-0300) | 1:1000 | 1:1000 | |
| GFAP | 2E1.E9 | Mouse IgG2b monoclonal | Biolegend (644702) | | | 1:5000 |
| GFP | | Chicken IgY polyclonal | Aves (1020) | | | 1:5000 |
| Ki67 | | Rabbit IgG polyclonal | Abcam (ab15580) | | 1:200 | |
| MAP2 | PCK-554P | Chicken IgY polyclonal | BioLegend (822501) | 1:1000 | 1:1000 | |
| Nestin | | Mouse IgG1 monoclonal | Abcam (ab6320) | 1:500 | 1:500 | |
| Neurofilament Marker (pan axonal, cocktail) | SMI-312 | Mouse IgG1 / IgM monoclonal | BioLegend (837904) | 1:1000 | 1:1000 | |
| Oct 4 | C-10 | Mouse IgG2b monoclonal | Santa Cruz (sc-5279) | 1:100 | | |
| PAX6 | Poly19013 | Rabbit IgG polyclonal | Biolegend (901301) | 1:200 | | |
| PSD95 | EP2652Y | Rabbit IgG monoclonal | Abcam (ab76115) | 1:200 | | |
| PSD95 | D27E11 | Rabbit IgG monoclonal | Cell Signaling (3450) | 1:400 | | |
| SATB2 | | Rabbit IgG polyclonal | Abcam (ab34735) | 1:200 | 1:200 | |
| SOX2 | D9B8N | Rabbit IgG monoclonal | Cell Signaling (23064S) | 1:200 | | 1:1000 |
| SOX2 | | Mouse IgG1 monoclonal | Santa Cruz (sc-365823) | 1:200 | 1:200 | |
| SSEA-4 | | Mouse IgG3 monoclonal | Santa Cruz (sc-21704) | 1:100 | | |
| SYNAPSIN-1 | | Rabbit IgG polyclonal | Abcam (ab64581) | 1:300 | | |
| SYNAPSIN-1 | D12G5 | Rabbit IgG monoclonal | Cell Signaling (5297) | 1:400 | | 1:5000 |
| TOMM20 | EPR15581-54 | Rabbit IgG monoclonal | Abcam (ab186735) | 1:250 | | |
| TRA 1-60 | | Mouse IgM monoclonal | Santa Cruz (sc-21705) | 1:100 | | |
| TRA 1-81 | | Mouse IgM monoclonal | Santa Cruz (sc-21706) | 1:100 | | |
| Tubulin β-3 (TUBB3) | Poly18020 | Rabbit IgG polyclonal | BioLegend (802001) | 1:1000 | 1:1000 | 1:5000 |
| VGAT | 117G4 | Mouse IgG3 monoclonal | Synaptic Systems (131 011) | 1:200 | | |
| VGLUT1 | | Rabbit IgG polyclonal | Synaptic Systems (135 302) | 1:200 | | |

**Supplementary Table 11. Secondary antibodies used in this study.**

| **Secondary antibody** | **Conjugate** | **Source** | **Dilution** |
|---|---|---|---|
| Goat anti-Rabbit IgG (H+L) | Alexa Fluor 633 | TermoFisher SCIENTIFIC (A-21070) | 1:500 |
| Goat anti-Rabbit IgG (H+L) | Alexa Fluor 546 | TermoFisher SCIENTIFIC (A-11010) | 1:1000 |
| Goat anti-Mouse IgG (H+L) | Alexa Fluor 488 | TermoFisher SCIENTIFIC (A-11001) | 1:1000 |
| Goat anti-Chicken IgY (H+L) | Alexa Fluor 633 | TermoFisher SCIENTIFIC (A-21103) | 1:500 |
| Goat anti-Rat IgG (H+L) | Alexa Fluor 555 | TermoFisher SCIENTIFIC (A-21434) | 1:1000 |
| Donkey anti-Mouse IgG (H+L) | Alexa Fluor 555 | TermoFisher SCIENTIFIC (A-31570) | 1:1000 |
| Donkey anti-Mouse IgG (H+L) | Alexa Fluor 647 | TermoFisher SCIENTIFIC (A-31571) | 1:1000 |
| Goat anti-Rabbit IgG H&L | HRP | Abcam (ab6721) | 1:200000 |
| Goat anti-Chicken IgY H&L | HRP | Abcam (ab6877) | 1:10000 |
| Rabbit anti Mouse IgG H&L | HRP | Abcam (ab6728) | 1:200000 |

**Supplementary Table 12. Primers used for qPCR experiments.**

| Genes | Forward primer 5'-3' | Reverse primer 5'-3' |
|---|---|---|
| *ACTB* | GGACTTCGAGCAAGAGATGG | AGCACTGTGTTGGCGTACAG |
| *GEPH* | TGTCACTCCAGAGGCCACAA | AGAGCATGCCCAGAGGTGTA |
| *MAP2* | CGCTGCGATCCCCTGATTTT | AGCTCTGCGACTAAGCAAGT |
| *MAP2* | CGAAGCGCCAATGGATTCC | TGAACTATCCTTGCAGACACCT |
| *PSD95* | AGAGACCAAGAGCTCCCAGG | CCTGGAAGAGGTCGCTATGC |
| *PSD95* | CCAGAGACCAAGAGCTCCCA | GCCTGGAAGAGGTCGCTATG |
| *SYN-1* | ACCCAGCCAGGACGTG | CAGAGACTGGGATTTGTTGAGC |
| *VGLUT1* | CTTCGCCATCTCTGGGTTCA | TTGTGCTTAGTCATGGCCCC |

### Supplementary methods

#### Generation of isogenic NSCs from iPSCs

Isogenic NSCs were generated from isogenic iPSCs following a previously published GIBCO protocol [1]. When the iPSCs reached 20% confluency, they were rinsed in DPBS to remove residual dead cells that had not attached to the surface, and the iPSCs culture medium was replaced with Neural Induction Medium (NIM) (Supplementary Table 3). That day was considered as day 0 of induction. The following day, the cells were examined under a microscope to confirm slight morphological changes toward NSCs. On that day, the old medium was also replaced with 2.5 mL of fresh NIM. The cells were monitored daily under a microscope, and the medium was changed every other day until day 7 (DIV7), with a gradual increase in medium volume up to 5 mL per well. At DIV7, the medium was completely removed, the cells were rinsed in DPBS, and 1 mL of accutase was added and the plate was incubated for 5 minutes at 37 °C until cell detachment was observed. DPBS at double the volume of Accutase was added to the cells to neutralize the enzyme and a single cell suspension was transferred to a sterile 50 mL tube after filtration through a 70 µm cell strainer (Cat. No. 431751). The cell suspension was then centrifuged for 4 minutes at 300 g at room temperature, after which the supernatant was removed, and the cell pellet was resuspended in 1 mL of Neural Expansion Medium (NEM). The composition of this medium is shown in Supplementary Table 4. Cells were counted and seeded onto Geltrex-coated 6-well plates at a density of $5x10^5$ cells per well in NEM supplemented with 5 µM ROCK inhibitor. The following day, the medium was completely replaced to remove the ROCK inhibitor. Cells were monitored daily, and the medium was fully changed every other day until the cells reached the desired confluency, after which they were passaged or cryopreserved.

#### Differentiation of isogenic neurons from NSCs

Isogenic neurons were differentiated from NSCs between passages P5 and P10. NSCs were rinsed in DPBS and subsequently incubated in 1mL of Accutase for 4 minutes at 37°C. NEM at double the volume of Accutase was added to the cells to neutralize the enzyme and a single cell suspension were generated by triturating. NSCs were seeded in 1 mL of NEM at a density of $2x10^5$ cells onto previously Geltrex-coated coverslips placed in 24-well plates. In addition to coverslips, cells were also seeded onto previously Geltrex-coated 6-well plates at a density of $1x10^6$ cells per well in 2 mL of NEM. That day was considered as day 0 of differentiation (DIV0). The following day, half of the NEM was removed and replaced with an equal volume of fresh BrainPhys medium (Supplementary Table 5). The medium was changed twice per week by removing half of the old medium and adding the same volume of fresh medium. Isogenic neurons were differentiated until predetermined time points (DIV14, DIV40, and DIV60). At these time points, neurons grown on coverslips were fixed and used for IF, while neurons cultured in 6-well plates were harvested using Accutase to achieve single cell suspension. Cells were centrifuged at 300 g for 4 minutes, the supernatant was removed, and the cell pellet was snap frozen on dry ice in sterile tubes and stored at -80 °C for WB and qPCR analyses.

**Generation of isogenic hCSAs**

Assembloids were generated following a previously published protocol [2], with the following modifications. Isogenic iPSCs were cultured in 6-well plates and E8 medium. When the cell colonies reached the desired confluency (70–80%), the medium was removed, and the cells were rinsed twice in DPBS and subsequently, 1 mL of Accutase was added, and the plate was incubated for 4 minutes at 37 °C. E8 Medium at double the volume of Accutase was added to the cells to neutralize the enzyme and a single cell suspension were generated by triturating. Cells were centrifuged for 4 minutes at 200 g, supernatant was removed and the cell pellet was resuspended in 1 mL of E8 medium supplemented with 10 μM ROCK inhibitor. Cells were counted and a total of 150 μL of the cell suspension was seeded into a 96-well clear round-bottom ultra-low attachment microplate at a density of $1x10^4$ cells per well. The plate was then centrifuged for 2 minutes at 200 g and placed in an incubator. That day was considered as day 0 of *in vitro* differentiation (DIV0). The following day, the cells were examined under a microscope to confirm spheroid formation, and the medium was not changed on that day. At DIV2, 120 μL of old medium was removed and replaced with 150 μL of fresh Essential 6 Medium (E6) (Supplementary Table 6). Immediately prior to the medium change, 2.5 μM Dorsomorphin and 10 μM SB-431542 were added to the E6 medium. That medium change was performed daily until DIV5. From DIV6 onward, the spheroids were directed toward either cortical (hCSs) or striatal (hStrSs) identity, with daily medium changes. The media composition used for striatal spheroids (Neural Differentiation Medium for hStrSs, NM hStrSs) and cortical spheroids (Neural Differentiation Medium for hCS, NM for hCSs) were shown in

Supplementary Tables 7 and 8. That patterning continued until DIV22, when a second set of factors were added to the differentiation medium. Daily medium changes were continued until DIV46, when cortical and striatal spheroids were fused to form cortico-striatal assembloids (hCSAs). To generate hCSAs, one cortical and one striatal spheroid were placed together at the bottom of a sterile 1.5 mL tube in 1 mL of Assembloid Differentiation Medium (ADM) (Supplementary Table 9). After two days, the medium was partially replaced by removing 750 μL of old medium and adding 1 mL of fresh medium. At DIV50, the formed assembloids were transferred to 24-well Clear Flat Bottom Ultra-Low Attachment Well Plates, and the differentiation medium was changed twice per week.

#### REFERENCES


1. Technologies, L. Induction of Neural Stem Cells from Human Pluripotent Stem Cells Using PSC Neural Induction Medium (MAN0008031 Rev A.0).
2. Miura Y, Li MY, Revah O, Yoon SJ, Narazaki G, Paşca SP. Engineering brain assembloids to interrogate human neural circuits. Nat Protoc. 2022; 17:15-35. https://www.nature.com/articles/s41596-021-00632-z.

**Supplementary Figures:**

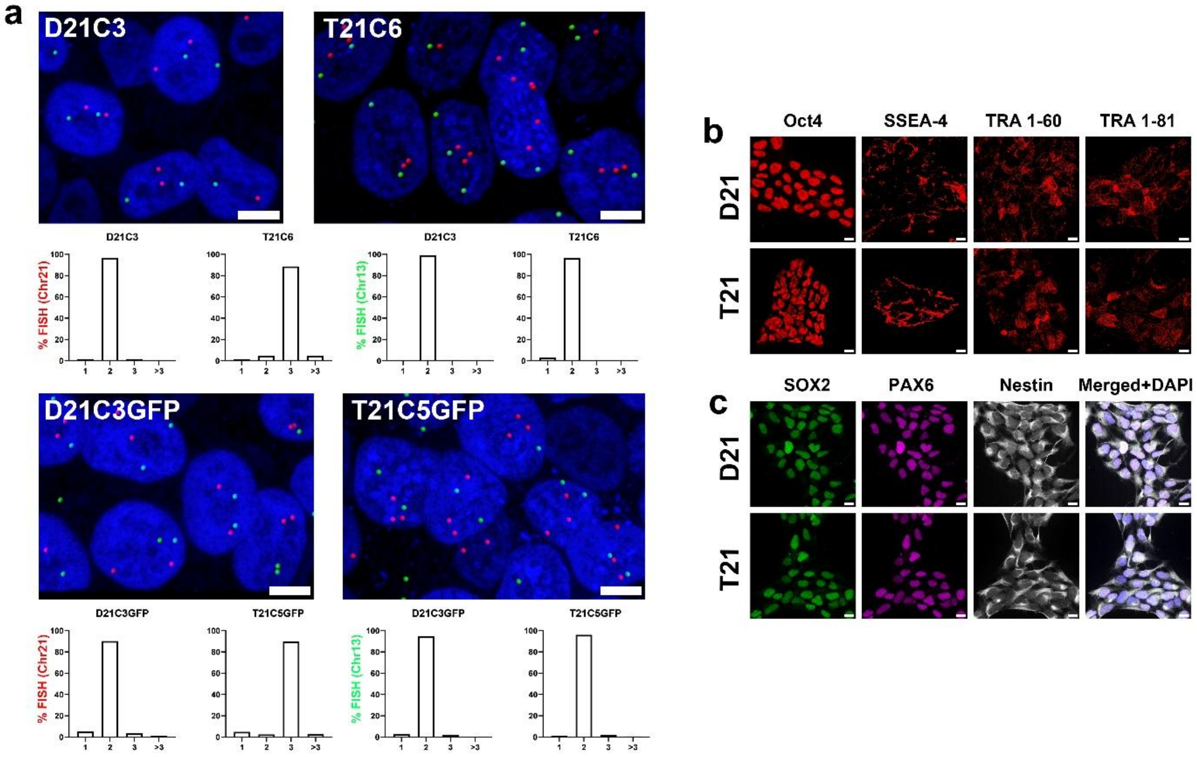


**Supplementary Fig. 1. Validation of isogenic iPSCs and NSCs**. **(a)** Representative confocal images of isogenic iPSCs used in this study, stained with FISH probe (green signal - chromosome 13, red signal - chromosome 21) and DAPI (blue). Scale bar: 10 µm. Our results showed that over 90% of D21 nuclei contained two chromosomes 21 (red signal), while over 90% of T21 nuclei contained three chromosomes 21 (red signal) and all clones contained two chromosomes 13 (green signal). The graphs were presented as percentage of nuclei with 1, 2, 3 and >3 signals. **(b)** Representative confocal images of isogenic iPSCs stained with following pluripotency markers: Oct4, SSEA-4, TRA 1-60 and TRA 1-81. Scale bar: 10 µm. Our data showed the same pattern of expression in both, D21 and T21 cells. **(c)** Representative images of isogenic NSCs stained with SOX2 (green), PAX6 (magenta), Nestin (white) and DAPI (blue). Scale bar: 10 µm. Our data showed the same pattern of expression in both, D21 and T21 cells.

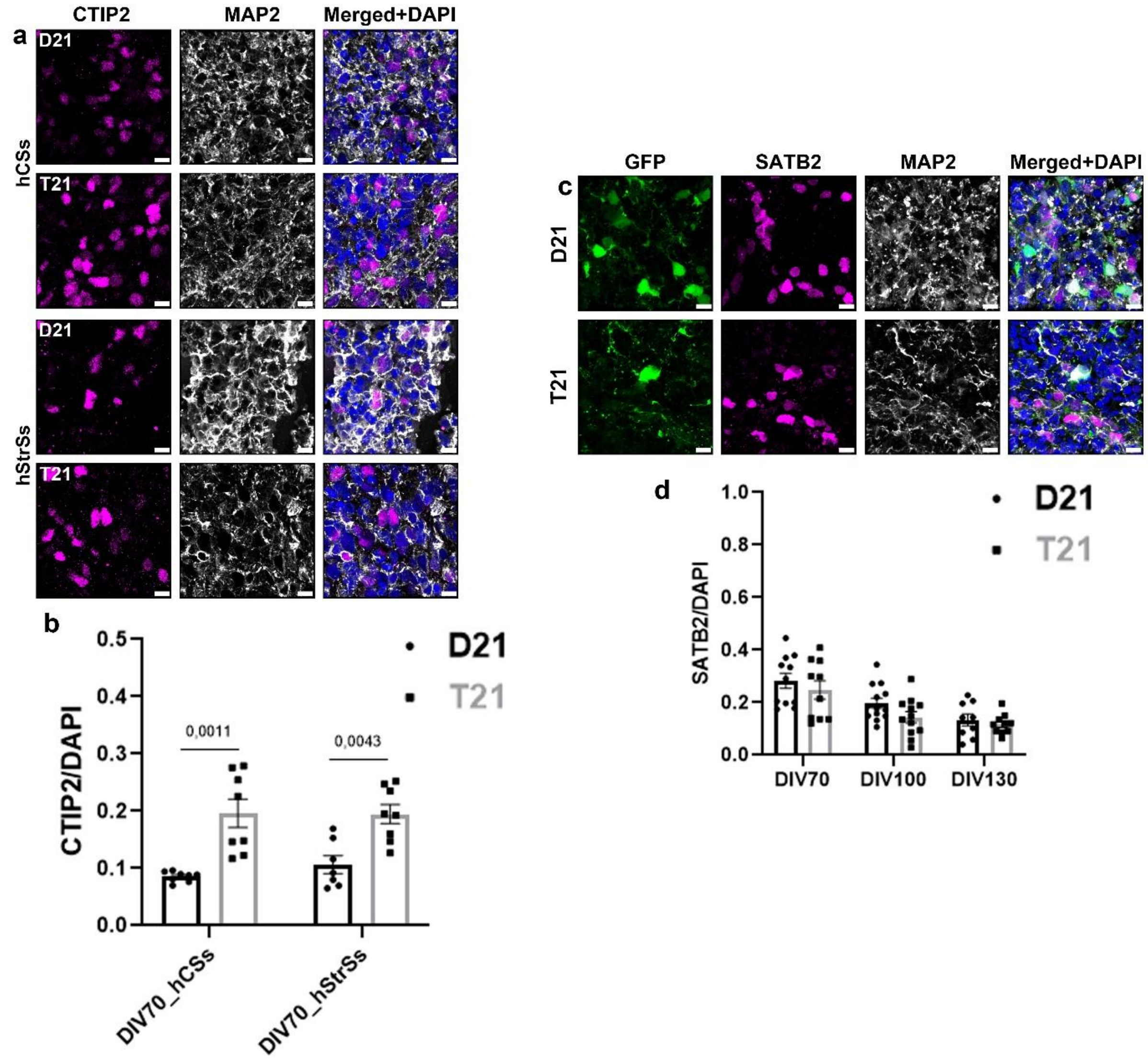


**Supplementary Fig. 2. Isogenic hCSAs express region-specific neuronal markers. (a)** Representative confocal images of isogenic hCSs and hStrSs stained with CTIP2 (magenta), MAP2 (white) and DAPI (blue) at DIV70. Scale bar: 10 µm. **(b)** T21 spheroids showed significantly higher expression of CTIP2 compared to D21 spheroids at DIV70. The y-axis represents expression of CTIP2 normalised by DAPI, while x-axis represents DIV 70. **(c)** Representative confocal images of isogenic hCSAs with GFP positive cells (green) stained with SATB2 (magenta), MAP2 (white) and DAPI (blue) at DIV100. Scale bar: 10 µm. **(d)** Our data showed the same level of expression throughout hCSAs *in vitro* differentiation. The y-axis represents expression of SATB2 normalised by DAPI, while x-axis represents DIV 70, DIV100 and DIV130. Graphs represent means ± SEM.

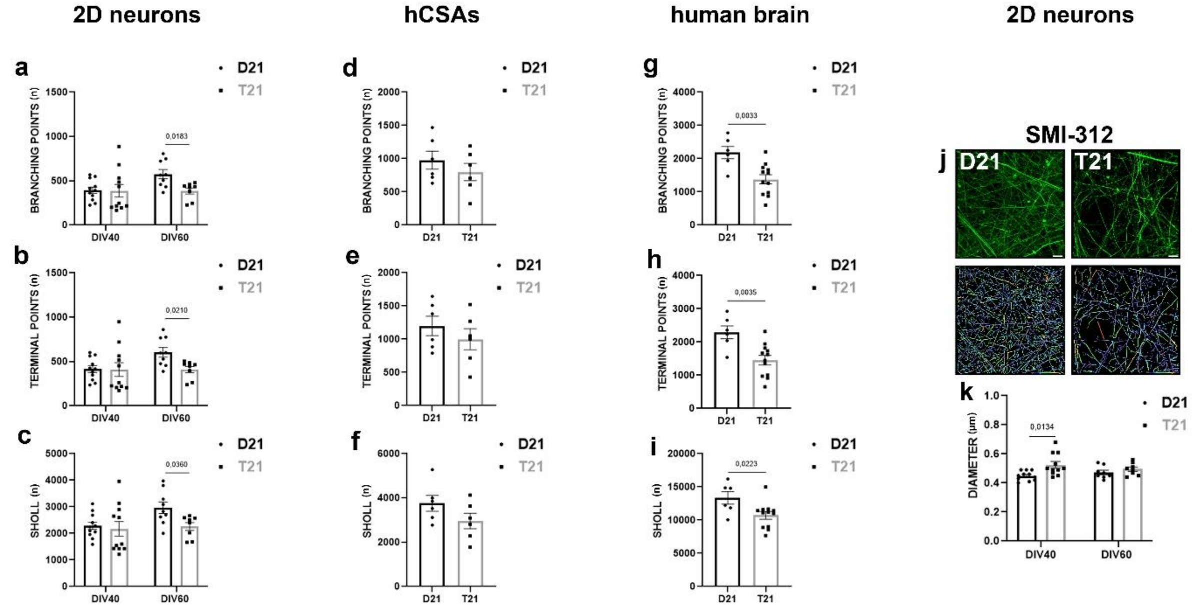


**Supplementary Fig. 3. T21 neurons showed abnormal neuronal morphology**. **(a-c)** 2D neurons: Isogenic (2D) MAP2 positive dendrites shown in main Fig. 3a. T21 dendrites showed fewer branching points, fewer terminal points and smaller Sholl diagram. **(d-f)** hCSAs: Isogenic (hCSAs) GFP/MAP2 positive dendrites shown in the main Fig. 3e. T21 dendrites showed fewer branching points, fewer terminal points and smaller Sholl diagram. **(g-i)** human brain: human cortical neurons shown in main Fig. 3j. T21 dendrites showed fewer branching points, fewer terminal points and smaller Sholl diagram. **(j-k)** 2D neurons: Representative confocal images of isogenic 2D axons and the same - Imaris 9.9.1 processed images. Different colour of neurites are Imaris software coded and represents diameter, length and branches. Scale bar: 10 µm. T21 axons showed larger diameter compared to the D21 axons. Branching point, terminal points and Sholl are presented as the number (n) of points per field of view, while the axonal diameter is presented in µm per field of view (y-axis). The x-axis represents

DIV40 and DIV60 for 2D neurons **(a-c, k)**, whereas for hCSAs **(d-f)** and human brain samples **(g-i)**, it represents genotype. Graphs represent means ± SEM.

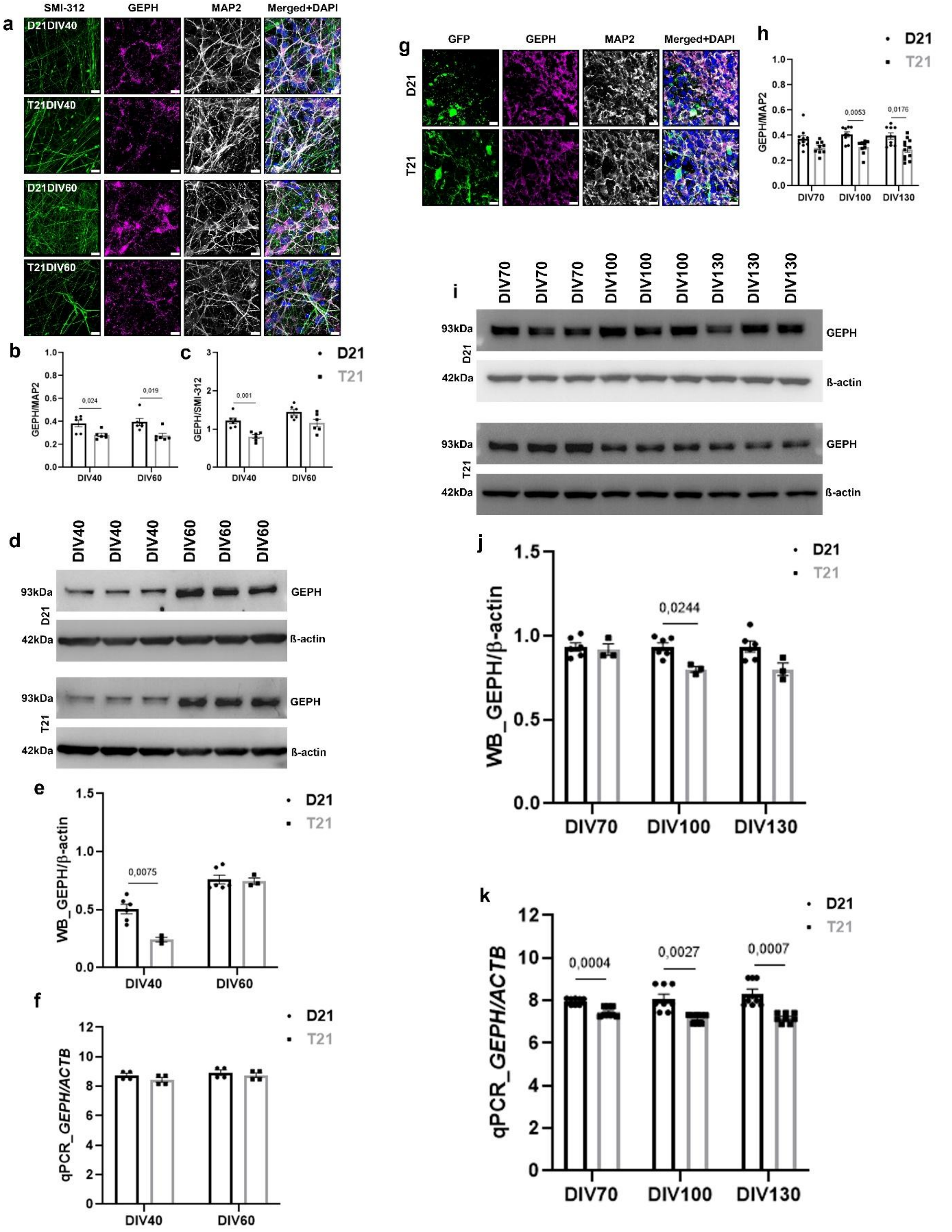

**Supplementary Fig. 4. Isogenic T21 neurons showed abnormal postsynaptic morphology.** **(a-f)** 2D neurons: **(a)** Representative confocal images of mature isogenic 2D neurons stained with SMI-312 (green), GEPH (magenta), MAP2 (white) and DAPI (blue) at DIV40 and DIV60. Scale bar: 10 µm. **(b)** T21 neurons showed significantly less GEPH positive synapses in MAP2 positive dendrites and **(c)** SMI-312 positive axons compared to D21 neurons throughout *in vitro* differentiation. The y-axis represents expression of GEPH normalised by MAP2 **(b)** and SMI-312 **(c)**, while x-axis represents DIV40 and DIV60. **(d)** Representative blots of GEPH/ß-actin throughout *in vitro* differentiation. **(e)** WB analysis of GEPH/ß-actin showed significantly lower expression in T21 neurons at DIV40. The y-axis represents expression of GEPH normalised by β-actin, while x-axis represents DIV40 and DIV60. **(f)** Analysis of *GEPH/ACTB* throughout *in vitro* differentiation showed the same level of gene expression in isogenic neurons. The y-axis represents expression of *GEPH* normalised by *ACTB*, while x-axis represents DIV40 and DIV60. **(g-k)** hCSAs: **(g)** Representative confocal images of isogenic hCSAs with GFP positive cells (green) stained with GEPH (magenta), MAP2 (white) and DAPI (blue) at DIV100. Scale bar: 10 µm. **(h)** T21 neurons showed significantly less GEPH positive synaptic puncta in MAP2 positive neurons compared to D21 neurons throughout *in vitro* differentiation. The y-axis represents expression of GEPH normalised by MAP2, while x-axis represents DIV70, DIV100 and DIV130. **(i)** Representative blots of GEPH/ß-actin throughout *in vitro* differentiation. **(j)** WB analysis of GEPH/ß-actin showed significantly lower expression in T21 hCSAs at DIV100. The y-axis represents expression of GEPH normalised by ß-actin, while x-axis represents DIV70, DIV100 and DIV130. **(k)** Analysis of *GEPH/ACTB* showed significantly lower gene expression in T21 hCSAs compared to D21 hCSAs throughout *in vitro* differentiation. The y-axis represents expression of *GEPH* normalised by *ACTB*, while x-axis represents DIV70, DIV100 and DIV130. Graphs represent means ± SEM.

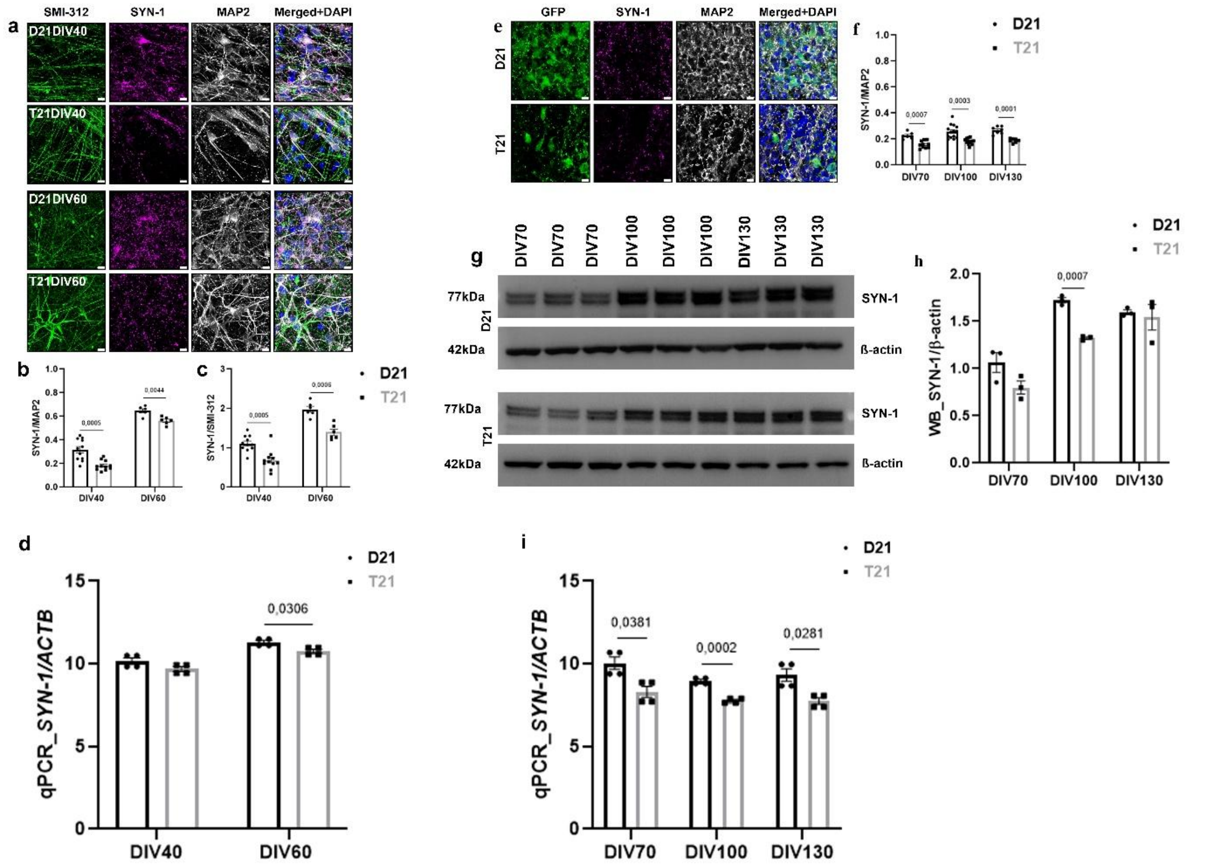


**Supplementary Fig. 5. Isogenic T21 neurons showed abnormal presynaptic morphology**. **(a-d)** 2D neurons: **(a)** Representative confocal images of mature isogenic 2D neurons stained with SMI-312 (green), SYN-1 (magenta), MAP2 (white) and DAPI (blue) at DIV40 and DIV60. Scale bar: 10 µm. **(b)** T21 neurons showed significantly less SYN-1 positive synapses in MAP2 positive dendrites and **(c)** SMI-312 positive axons compared to D21 neurons throughout *in vitro* differentiation. The y-axis represents expression of SYN-1 normalised by MAP2 **(b)** and SMI-312 **(c)**, while x-axis represents DIV40 and DIV60. **(d)** Analysis of *SYN-1/ACTB* showed significantly lower gene expression in T21 neurons at DIV60. The y-axis represents expression of *SYN-1* normalised by *ACTB*, while x-axis represents DIV40 and DIV60. **(e-i)** hCSAs: **(e)** Representative confocal images of isogenic hCSAs with GFP positive cells (green) stained with SYN-1 (magenta), MAP2 (white) and DAPI (blue) at DIV100. Scale bar: 10 µm. **(f)** T21 neurons showed significantly less SYN-1 positive synapses in MAP2 positive neurons compared to D21 neurons throughout *in vitro* differentiation. **(g)**

Representative blots of SYN-1/ß-actin throughout *in vitro* differentiation. **(h)** WB analysis of SYN-1/ß-actin showed significantly lower expression in T21 hCSAs at DIV100. The y-axis represents expression of SYN-1 normalised by ß-actin, while x-axis represents DIV70, DIV100 and DIV130. **(i)** Analysis of *SYN-1/ACTB* showed significantly lower gene expression in T21 hCSAs compared to D21 hCSAs throughout *in vitro* differentiation. The y-axis represents expression of *SYN-1* normalised by *ACTB*, while x-axis represents DIV70, DIV100 and DIV130. Graphs represent means ± SEM.

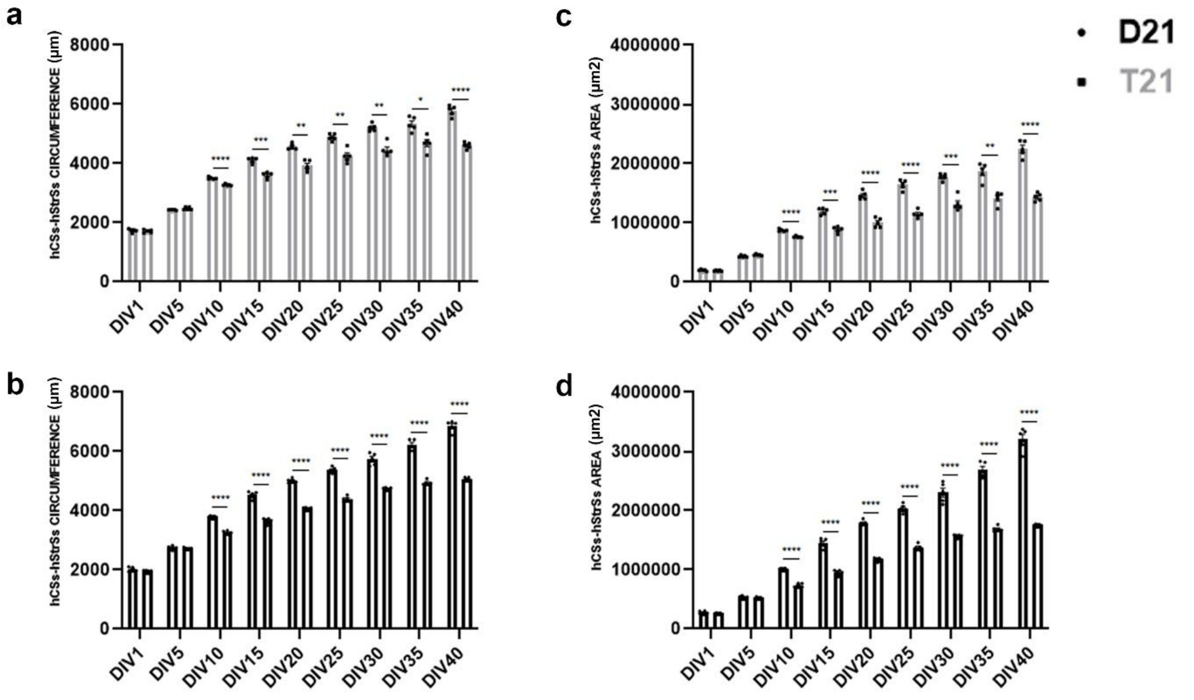


**Supplementary Fig. 6. T21 spheroids were significantly smaller compared to D21 spheroids**. **(a)** Analysis of the T21 hCSs and hStrSs circumference showed significantly higher values in hCSs throughout *in vitro* differentiation. **(b)** Analysis of the D21 hCSs and hStrSs circumference showed significantly higher values in hCSs throughout *in vitro* differentiation. **(c)** Analysis of the T21 hCSs and hStrSs area showed significantly higher values in hCSs throughout *in vitro* differentiation. **(d)** Analysis of the D21 hCSs and hStrSs area showed significantly higher values in hCSs throughout *in vitro* differentiation. Circumference **(a, b)** is presented in μm (y-axis), while area **(c, d)** is presented in $\mu m^2$ (y-axis). The x-axis represents DIV1-40 (a-d). The left column in each DIV represents hCSs, while the right column represents hStrSs (a-d). Graphs represent means ± SEM.

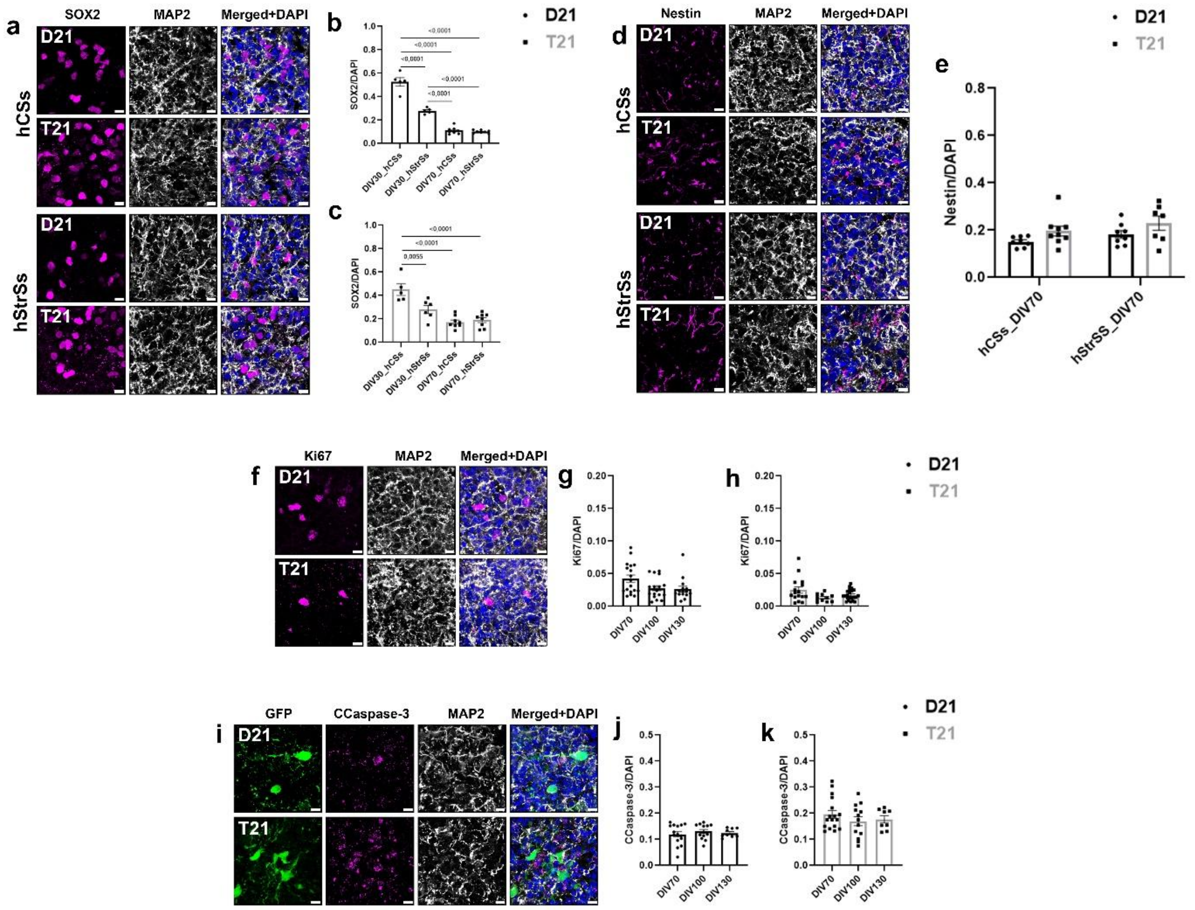


**Supplementary Fig. 7. Proliferation and cell death throughout *in vitro* differentiation**. **(a)** Representative confocal images of isogenic hCSs and hStrSs stained with SOX2 (magenta), MAP2 (white) and DAPI (blue) at DIV70. Scale bar: 10 μm. **(b)** D21 and **(c)** T21 spheroids showed significant decrease of SOX2 throughout *in vitro* differentiation. The y-axis represents expression of SOX2 normalised by DAPI, while x-axis represents DIV30 and DIV70 of **(b)** D21 and **(c)** T21 spheroids. **(d)** Representative confocal images of isogenic hCSs and hStrSs stained with Nestin (magenta), MAP2 (white) and DAPI (blue) at DIV70. Scale bar: 10 μm. **(e)** Our data showed the same level of Nestin expression at DIV70. The y-axis represents expression of Nestin normalised by DAPI, while x-axis represents DIV70. **(f)** Representative confocal images of isogenic hCSAs stained with Ki67 (magenta), MAP2 (white) and DAPI (blue) at DIV100. Scale bar: 10 μm. **(g-h)** Our data showed significantly fewer proliferating cells labelled with Ki67 in T21 throughout *in vitro* differentiation. The y-axis represents expression of Ki67 normalised by DAPI, while x-axis represents DIV70, DIV100 and DIV130 of **(g)** D21 and **(h)** T21 hCSAs. **(i)** Representative confocal images of isogenic hCSAs with GFP positive cells (green) stained with CCaspase-3 (magenta), MAP2 (white) and DAPI (blue) at DIV100. Scale bar: 10 μm. **(j-k)** Our data showed significantly more dead cells labelled with CCaspase-3 in T21 throughout *in vitro* differentiation. The y-axis represents expression of CCaspase-3 normalised by DAPI, while x-axis represents DIV70, DIV100 and DIV130 of **(j)** D21 and **(k)** T21 hCSAs. Graphs represent means ± SEM.
